# Hippocampal and thalamic afferents form distinct synaptic microcircuits in the mouse frontal cortex

**DOI:** 10.1101/2021.03.12.435140

**Authors:** Kourtney Graham, Nelson Spruston, Erik B. Bloss

**Affiliations:** Howard Hughes Medical Institute, Janelia Research Campus, Ashburn, VA 20147, USA; The Jackson Laboratory, 600 Main Street, Bar Harbor, ME 04609, USA

## Abstract

Neural circuits within the frontal cortex support the flexible selection of goal-directed behaviors by integrating input from brain regions associated with sensory, emotional, episodic, and semantic memory functions. From a connectomics perspective, determining how these disparate afferent inputs target their synapses to specific cell types in the frontal cortex may prove crucial in understanding circuit-level information processing. Here, we used monosynaptic retrograde rabies mapping to examine the distribution of afferent neurons targeting four distinct classes of local inhibitory interneurons and four distinct classes of excitatory projection neurons in mouse infralimbic cortex. Interneurons expressing parvalbumin, somatostatin, or vasoactive intestinal peptide received a large proportion of inputs from hippocampal regions, while interneurons expressing neuron-derived neurotrophic factor received a large proportion of inputs from thalamic regions. A more moderate hippocampal-thalamic dichotomy was found among the inputs targeting excitatory neurons that project to the basolateral amygdala, lateral entorhinal cortex, nucleus reuniens of the thalamus, and the periaqueductal gray. Together, these results show a prominent bias among hippocampal and thalamic afferent systems in their targeting to genetically or anatomically defined sets of frontal cortical neurons. Moreover, they suggest the presence of two distinct local microcircuits that control how different inputs govern frontal cortical information processing.

## Introduction

Neuronal circuits in the frontal cortex mediate some of the mammalian brain’s most advanced forms of cognition, including the context-dependent selection of goal-directed behaviors (Miller, 2000). Logically, this function requires information relevant to the results from previous experiences (Hasegawa et al., 2000) as well as highly processed sensory information reflecting potentially new and relevant contextual cues. Electrical recordings from frontal cortex in awake, behaving animals have shown that individual frontal cortical neurons encode diverse representations of behaviorally relevant features, including sensory features, spatial locations, task temporal structure, and cues that predict previously rewarded or non-rewarded outcomes. Although the presence of these mixed representations in frontal cortex is intriguing (Hirokawa et al., 2019; Kennerley and Wallis, 2009; Machens et al., 2010; Rigotti et al., 2013), it remains unclear how such signals are generated.

These mixed representations could result from a neuroanatomical organization in which specific afferent information streams are connected to specific subsets of postsynaptic frontal cortical neurons. In this scenario, cell-type specific forms of synaptic connectivity would provide a hardwired constraint on the possible representations that could be signaled by specific frontal cortical neurons. Such a scenario would be consistent with previous connectomics results demonstrating both cell-type and subcellular specificity and precision within circuit synaptic architectures (Bloss et al., 2018; Druckmann et al., 2014; Kasthuri et al., 2015). Conversely, all frontal cortical neurons might receive input from each afferent pathway yet produce mixed representations through cell-autonomous forms of synaptic plasticity or task-specific forms of neuromodulation. We sought to test which of these scenarios predominated on inhibitory and excitatory neurons in the mouse frontal cortex.

Generating brain-wide maps of connected neurons has been a major challenge for neuroscience given the submicrometer scale of synapses connecting two neurons yet the 100- to 1000-fold greater scale of axonal and dendritic processes within the brain (Lichtman and Denk, 2011). To circumvent the need for ultrastructural visualization of synaptic connections, the transsynaptic and retrograde transport properties of rabies viruses have been exploited to produce maps of connected neurons across long distances at cellular resolution (Luo et al., 2018; Ugolini, 2011; Wall et al., 2010; Wickersham et al., 2007). Moreover, the recent ability to deliver rabies-derived reagents to molecularly- or anatomically- defined cell types offer the potential to determine whether different cell types have distinct sets of inputs and outputs across the entire brain.

The extent to which rabies experiments can produce accurate maps of connected neurons is dependent on the efficiency of the transsynaptic retrograde transport. Nearly all cell-type specific rabies mapping experiments have used a single genetically modified strain of rabies virus (SADΔG); recently, Reardon and colleagues (Reardon et al., 2016) have shown that the CVS strain of rabies permits significantly greater labeling of long-range connected circuits. Here, we used the CVSΔG variant to perform cell-type-specific, monosynaptic, retrograde rabies tracing to determine the input patterns to specific cell classes in the infralimbic (IL) region of the mouse frontal cortex. Using targeted knockin Cre driver lines, we mapped the brain-wide pattern of afferent neurons forming synapses on IL interneurons expressing parvalbumin (PV), somatostatin (SST), vasoactive intestinal peptide (VIP), and neuron-derived neurotrophic factor (NDNF). Using a AAVRetro-Cre strategy to gain genetic access to projection neurons (Tervo et al., 2016), we also mapped the brain-wide pattern of afferent neurons targeting specific IL excitatory projection neurons. We chose projection neurons targeting the basolateral amygdala (BLA), lateral entorhinal cortex (LEC), nucleus Reuniens of the thalamus (RE), or periaqueductal gray (PAG) because they have been implicated in IL-dependent control of fear-related behaviors (Bloodgood et al., 2018; Ramanathan et al., 2018; Rozeske et al., 2018; Xu et al., 2012).

Our results support a model in which different IL neurons receive overlapping sets of afferent inputs, yet the proportion of each pathway varied across the cell classes. Specifically, we show that IL PV-Cre, SST-Cre, and VIP-Cre interneurons each receive the largest proportion of their afferent inputs from hippocampal area CA1, while the largest proportion of inputs to NDNF-Cre interneurons originate from RE. A similar dichotomy in the receipt of hippocampal and thalamic inputs remained evident across excitatory projection neurons. Because the interneurons examined here provide functionally distinct forms of inhibition targeted to different dendritic regions of IL projection neurons, our data suggest the presence of two distinct spatially organized microcircuits that govern information transforms in IL projection neurons. Consistent with the notion that CVSΔG provides higher transsynaptic efficiency at synapses forming long-range pathways, our results differ substantially from published work using the SADΔG virus to map inputs to neurons in mouse frontal cortex (Ahrlund-Richter et al., 2019; Sun et al., 2019). Finally, we have created a public resource to accompany this work, <u>r</u>abies-<u>a</u>ssisted <u>i</u>nterrogation of <u>s</u>ynaptic <u>i</u>nfralimbic <u>n</u>etworks (or RAISIN; http://raisin.janelia.org), that provides analysis and visualization of these datasets so future experiments can relate the underlying structural features of frontal cortical circuits to their functional computations.

## Results

### Maps of IL inputs and outputs

To determine the diversity of cortical and subcortical inputs to and outputs from IL, we first made input/output maps from mouse IL cortex by direct injection of recombinant adeno-associated viruses (rAAVs) expressing fluorescent proteins that travel in a retrograde (i.e., RetroAAV (Tervo et al., 2016)) or anterograde (i.e., rAAV2/1) manner and compared these results to those obtained from fluorescent conjugated tracers (CTB-555 and WGA-555). Major inputs to the mouse IL cortex arise from area CA1 of the hippocampus, mediodorsal (Md-Th) and RE, the BLA, and from neighboring regions within frontal cortex, all consistent with data from rat (Hoover and Vertes, 2007) and nonhuman primate (Barbas, 2000; Ongur and Price, 2000). Conversely, the major IL outputs are neighboring regions of the frontal cortex, the dorsomedial and ventral striatal regions, both Md-Th and RE, the lateral hypothalamus, and the BLA. IL circuits are thus organized as efferent-only pathways (e.g., IL-to-ventral striatum with no ventral striatum-to-IL connection), afferent-only pathways (e.g., CA1-to-IL projections with no IL-to-CA1 projection), or as reciprocal loops (e.g., connections to and from Md-Th, RE, and BLA) (**Supplementary Figure 1**).

### Genetic and viral access to non-overlapping cell classes in IL cortex

These coarse input/output maps are useful insofar as they constrain the possible routes of information flow through mouse IL cortex. However, such maps lack the ability to discern whether afferent regions projecting to IL use similar or different patterns of connectivity onto specific postsynaptic cell types. Circuit mapping strategies that take advantage of the transsynaptic spread of rabies virus have been developed to answer precisely such questions. For these strategies to generate interpretable maps, however, they must be employed to distinct postsynaptic cell classes that have little to no minimal overlap. We used two different strategies to gain genetic access to distinct, nonoverlapping IL neurons: in the first, we used transgenic targeted knockin mice where Cre recombinase was driven by the promoter of genes expressed in subsets of cortical interneurons. In the second, we used intracranial injections of RetroAAV-Cre in downstream IL target regions to drive Cre recombinase in IL projection neurons.

We first confirmed that PV-Cre (Hippenmeyer et al., 2004), VIP-Cre (Taniguchi et al., 2011), SST-Cre (Taniguchi et al., 2011), and NDNF-Cre (Tasic et al., 2016) driver lines permit genetic access to distinct sets of inhibitory interneurons in IL. Using triple-label *in situ* hybridization and confocal microscopy, we found that Cre expression within each driver line demonstrated high specificity and efficiency, and that overlap between Cre-expressing neurons and marker genes for the other driver lines was low (< 1% in all comparisons for all lines) (**Figure 1A-C**). PV-Cre, VIP-Cre, SST-Cre, and NDNF-Cre neurons differed in terms of the laminar location within IL (**Supplementary Figure 2**), and reconstruction of virus-labeled neurons from these Cre driver lines demonstrated large differences in their dendritic morphology (**Figure 1C and Supplementary Figure 2**).

**Figure 1.**
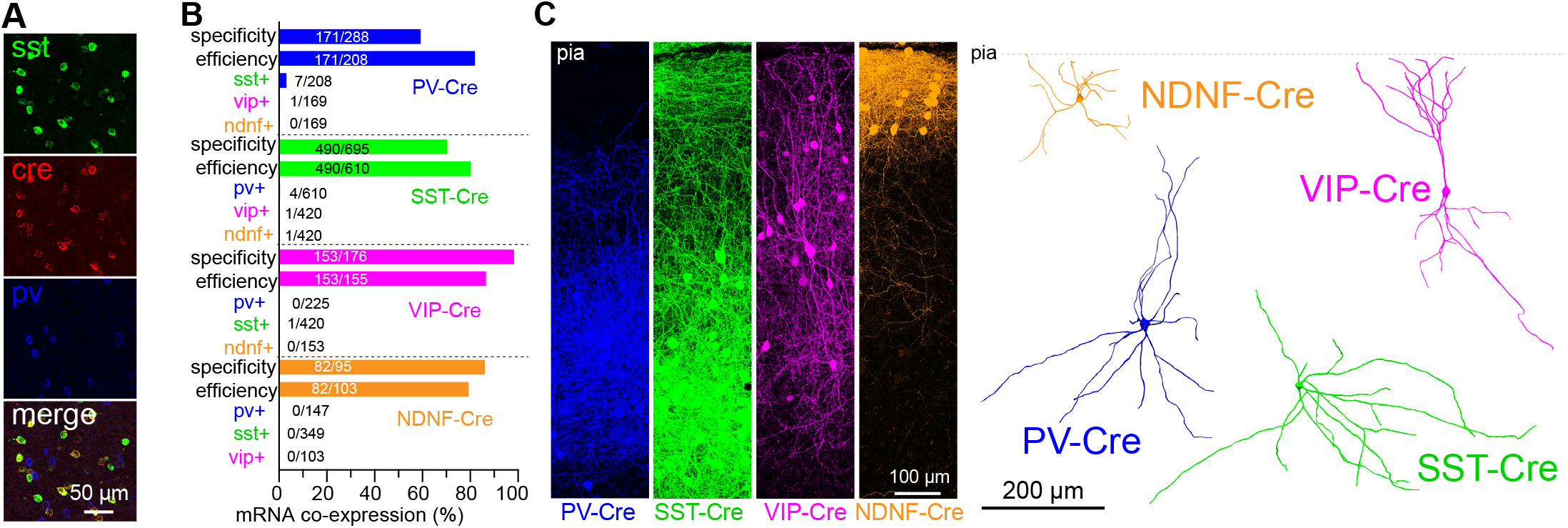
Genetic access to distinct interneuron classes in IL cortex. **A)** Example of a triple-label fluorescence *in situ* hybridization experiment from an SST-Cre mouse with probes labeling *sst*, *cre*, or *pv*. **B)** Quantification of the specificity, efficiency, and marker gener expression overlap across each Cre driver line in IL cortex. **C)** Transduction of these driver lines with a viral Cre-dependent GFP reporter (left; lines are pseudocolored) permitted visualization and reconstruction of neuronal morphology (right); see **Supplementary Figure 2** and **Table S6** for the quantification of soma locations and dendritic morphologies.

In the absence of Cre driver lines for different projection neurons, we gained genetic access via axonal transduction of a new designer AAVRetro virus optimized for retrograde transport (Tervo et al., 2016) (e.g., in a manner similar in spirit to (Schwarz et al., 2015)). To determine the overlap between IL projection neurons, we visualized pairs of projection neurons using a viral dual recombinase approach in which AAVRetro-Cre and AAVRetro-FlpO were injected into separate downstream targets and labeled in IL with Cre- and FlpO-dependent viral fluorescent reporters. We found that all four sets of projection neurons were spatially intermingled in IL but arose from largely non-overlapping populations of superficial (e.g., BLA-targeting and LEC-targeting) or deep-layer (PAG-targeting and RE-targeting) pyramidal cells (**Figure 2A-B**), consistent with previous reports from the adjacent dorsal PL region (Cheriyan et al., 2016; Collins et al., 2018; Little and Carter, 2013). Reconstructions of projection neurons demonstrated nearly identical dendritic morphologies within each superficial or deep layer pair (**Supplementary Figure 2**), permitting a stringent comparison to be made across separate projection neurons with similar dendritic patterns. Thus, interneuron Cre driver lines and RetroAAV-Cre transduction of projection neurons permit genetic access to non-overlapping cell classes within the mouse IL cortex.

**Figure 2.**
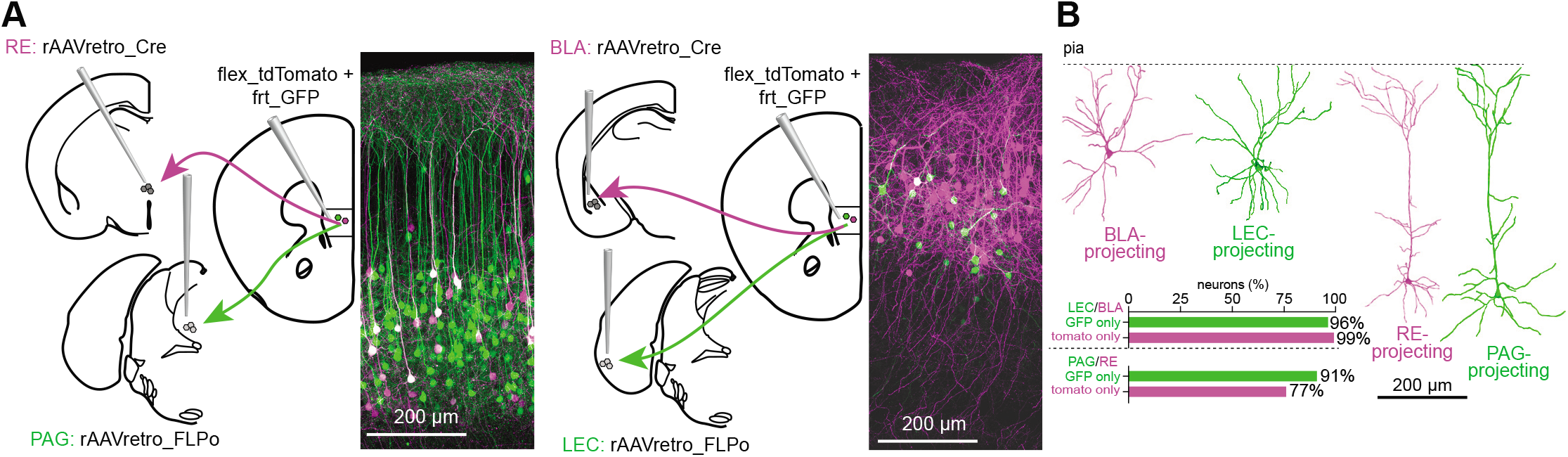
A retrograde viral approach permits access to distinct excitatory projection neurons in IL cortex. **A)** Schematic showing labeling of IL projection neurons using a dual Cre and FLPo retrograde viral recombinase strategy. **B)** Reconstructions of the dendritic morphology of BLA-, LEC-, RE-, and PAG-projecting neurons and their cellular overlap; see **Supplementary Figure 2 and Table S6** for the quantification of soma locations and dendritic morphologies.

### Quantification of RabV starter cells

To transduce Cre-expressing neurons with CVS-N2cΔG EGFP(EnvA) RabV (Reardon et al., 2016), two Cre-dependent AAVs were injected into IL: one encoding the TVA receptor required for entry of the modified RabV, and the other encoding the N2c-ΔG protein required for RabV transsynaptic movement. Both of these AAVs also expressed the far-red fluorophore mKate2, permitting the identification of the neurons competent for subsequent RabV transduction and transsynaptic spread (i.e., “starter cells”). Neurons transduced by helper viruses *and* RabV express both GFP and mKate2, while RabV-labeled presynaptic neurons express GFP only (**Figure 3A-C**). To quantify the number of mKate2+ or GFP+ cells, we developed a semiautomated analysis pipeline that aligns, thresholds, assigns, and counts GFP+ neurons in individual brain regions. We first validated this pipeline by control experiments that revealed a low rate of false positive or false negatives compared to manual neuron counts (**Supplementary Figure 3**). We quantified and assigned the number of mKate2^+^/GFP^+^ double-labeled “starter cells” near the injection sites and found these starter cells enriched within IL cortex in both interneuron classes and projection neuron experiments (**Figure 3D**). The total number of starter cells (summed across all regions) correlated strongly with the numbers of presynaptic neurons labeled by RabV-GFP (interneurons: n=12, Spearman r= 0.83, p=0.0015; virus-labeled projection neurons: n=13, Spearman r= 0.79, p=0.002) (**Figure 3E**).

**Figure 3.**
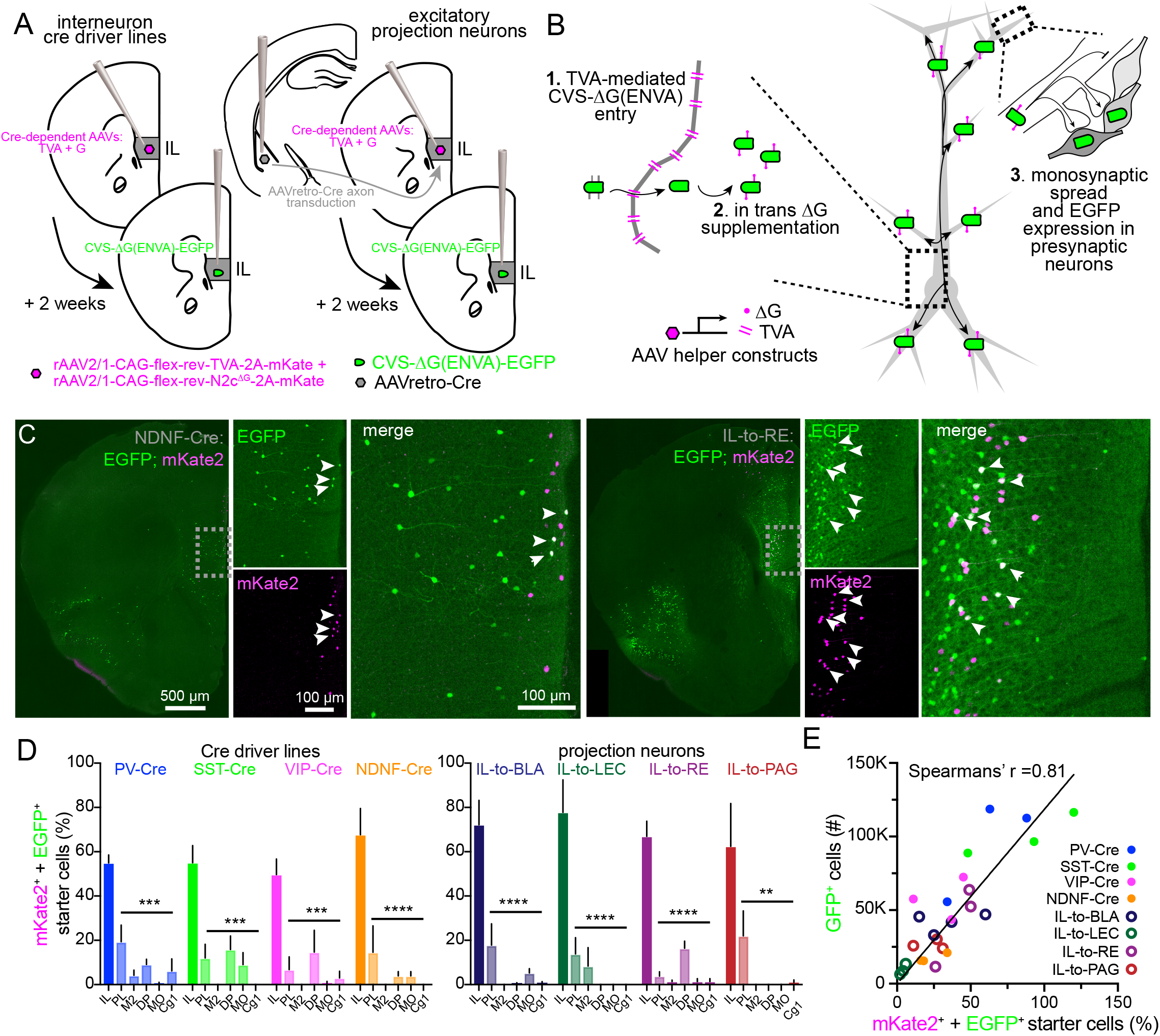
Labeling afferents to specific neuronal classes. A, The injection configuration used to transduce RabV-competent starter cells in Cre driver lines or projection neuron classes. B, in this approach, TVA and ΔG are expressed via Cre-dependent rAAVs to render specific neurons competent to take up and transport the CVS ΔG(ENVA)-EGFP. C, ‘Starter cells’ at the injection site were identified by the dual expression of both mKate2 and EGFP); the left four images are from an experiment with IL NDNF-Cre neurons (note the localization of starter cells to layer I) and the right four are from an experiment with IL-to-RE neurons (note the localization of starter cells to layer V). D, Starter cells were preferentially localized within IL relative to the neighboring cortical regions. E, Starter cell numbers correlated strongly with the total number of EGFP-labeled presynaptic neurons. (Note one IL-to-BLA mouse was excluded from the starter cell analyses on accound of poor mKate2 signal). See Supplementary Figure 3 and Table S2 for additional information and statistical analyses.

To determine the viral ‘leak’ in our CVS-N2cΔG EGFP(EnvA) approach, we performed identical experiments in the absence of any transgenic or viral Cre expression (e.g., in WT mice). Since cortical neurons should not be competent to express the Cre-dependent helper constructs that were injected in IL, there should be very few GFP-expressing neurons. Consistent with this notion, these experiments yielded 488 ± 118 GFP^+^ neurons (mean ± SEM, n=3 mice) scattered across the mouse brain, suggesting a small but nonzero leak of our viral system (similar to (Miyamichi et al., 2013; Weissbourd et al., 2014)) (**Supplementary Figure 3**). By contrast, the experiments with transgenic or viral Cre expression yielded 49,024 ± 6,698 GFP^+^ labeled neurons (mean ± SEM, n=26 mice), (**Supplementary Figure 3**), suggesting that this leak contributes approximately 1% of the total labeled neurons in an experimental sample.

### Distribution of afferent neurons targeting IL inhibitory and excitatory neurons

We examined the spatial distribution of the GFP^+^ input neurons along the anterior-posterior (A-P) axis of the brain from our inhibitory interneuron and excitatory projection neuron datasets. When comparing the pooled inhibitory datasets to the pooled excitatory neuron datasets, the input fractions along the anterior-posterior axis were statistically indistinguishable (**Figure 4A,B and Supplementary Figure 4**). Both datasets featured two prominent and well-separated peaks along the A-P axis: one that lies between 0.5 mm and 1.5 mm posterior to bregma (the “anterior peak”), and a second that lies between 3.5 to 4.0 mm posterior to bregma (the “posterior peak”). These datasets suggest that the majority of presynaptic neurons targeting IL neuronal classes labeled by CVS-N2c-ΔG EGFP(EnvA) RabV are long-range projection neurons (i.e., arise from outside local frontal cortical regions). This result differs dramatically from previous IL or PL RabV experiments that concluded inputs to both inhibitory and excitatory neurons were predominantly made *within* frontal cortical circuitry (Ahrlund-Richter et al., 2019; Sun et al., 2019).

**Figure 4.**
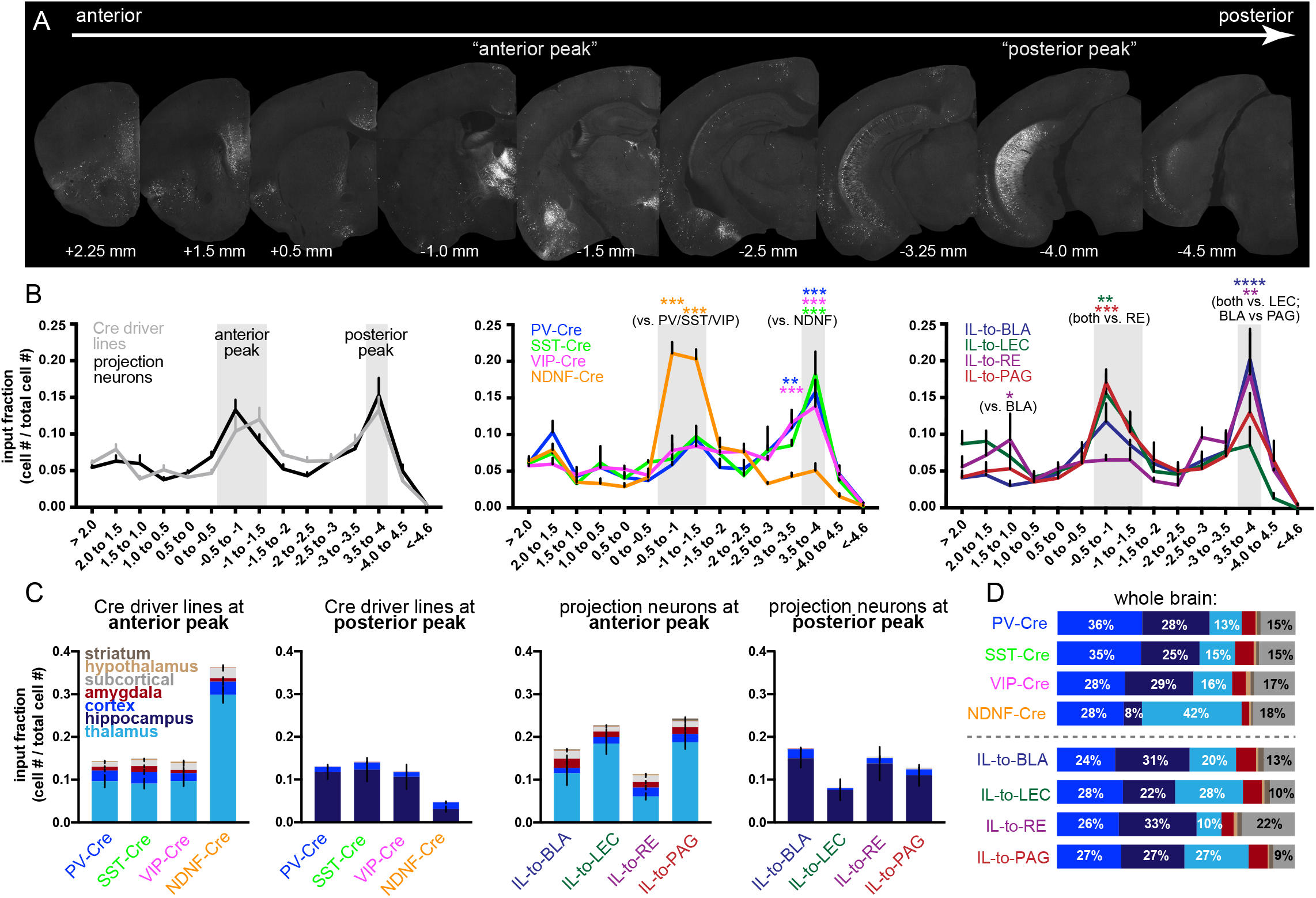
Spatial distribution of afferent neurons targeting different IL cell types. A, Images from an IL-to-BLA sample showing presynaptic cells along the anterior-posterior axis (top image panels) with high density of neurons at −1.0 mm and −1.5 mm (referred to as the “anterior peak”), as well as at −4.0 mm (the “posterior peak”). B, Input fraction of EGFP^+^neurons along the A-P axis between the pooled inhibitory Cre driver lines and the excitatory projection classes (left), among the inhibitory interneuron Cre driver lines (center) and among the excitatory projection neurons (right). C, Neurons assigned to the thalamus, hippocampus, cortex, amygdala, subcortical, hypothalamus or striatum at the anterior and posterior peaks in the inhibitory Cre driver lines and projection neurons. D, Normalized percentage of inputs from these groups aross the whole brain. See Supplementary Figure 4 and Table S3 for additional information and statistical details.

When we examined these distributions across the different inhibitory Cre driver lines, we found that PV-Cre, SST-Cre, and VIP-Cre neurons had virtually identical patterns of input fraction of presynaptic neurons along the A-P axis. The largest input fraction to these three interneurons was found at the posterior peak; in contrast, the largest proportion of input to NDNF-Cre neurons was found at the anterior peak (**Figure 4B and Supplementary Figure 4**). At both peaks, the magnitude of the effect was roughly two-fold (i.e., the input fraction was roughly two-fold larger for NDNF-Cre mice compared to PV-Cre/SST-Cre/VIP-Cre at the anterior peak and PV-Cre/SST-Cre/VIP-Cre neurons had a two-fold greater input fraction at the posterior peak; p<0.0001 for each comparison) and was evident in each replicate sample (**Supplementary Figure 4**). Among the excitatory projection classes, differences were also evident in the input fractions at the anterior-posterior axis peaks. Specifically, LEC- and PAG-projecting neurons had a larger input fraction at the anterior peak compared to REprojecting neurons (p<0.005 and p<0.0005, respectively), while LEC-projecting neurons had a smaller input fraction at the posterior peak compared to BLA-projecting and RE-projecting neurons (p<0.0001 and p<0.005) (**Figure 4B and Supplementary Figure 4**).

We next examined how the input fractions within each peak related to brain structures (**Figure 4C**). At the anterior peak, all interneurons received the majority of inputs from the thalamus, yet NDNF-Cre neurons received roughly 3-fold greater proportion of thalamic inputs compared to PV-Cre, SST-Cre, or VIP-Cre neurons. Conversely, at the posterior peak, all interneurons received the majority of inputs from the hippocampus yet PV-Cre, SST-Cre, and VIP-Cre received approximately 3-fold greater inputs from hippocampal regions than NDNF-Cre neurons. Similar patterns were found among the projection neurons: LEC-projecting and PAG-projecting neurons received roughly 2-to-3-fold greater proportion of inputs from the thalamus than BLA- or RE-projecting neurons. Conversely, BLA- and RE-projecting neurons received approximately 2-fold greater inputs from hippocampal regions than LEC-projecting neurons. The differential receipt of inputs from neurons across brain structures at each peak was recapitulated when we considered the overall proportion of input neurons across the brain (**Figure 4D**). Collectively, these results show that IL neuron classes receive different proportions of inputs from neurons in the thalamus and hippocampus.

### Regional differences in the proportion of afferent neurons targeting IL interneurons

We next considered the distribution of presynaptic neurons across functionally relevant groups of thalamic and cortical nuclei. We used a recently described taxonomy of thalamic regions (Phillips et al., 2019) to determine whether inputs from major thalamic nuclei differentially targeted IL interneurons. We found that regions in the secondary thalamic group (e.g., Md-Th, AM, VM, VA) and RE (which forms its own distinct molecular class within the thalamus) each had a greater input fraction to NDNF-Cre neurons than neurons in the other three Cre driver lines (**Figure 5A**). Using information modality to parse cortical regions into coarse functional groups (e.g., frontal association, motor, somatosensory, visual, etc.), we found that both PV and SST-Cre mice had greater input fractions from frontal association regions compared to VIP or NDNF-Cre mice (**Figure 5A**).

**Figure 5.**
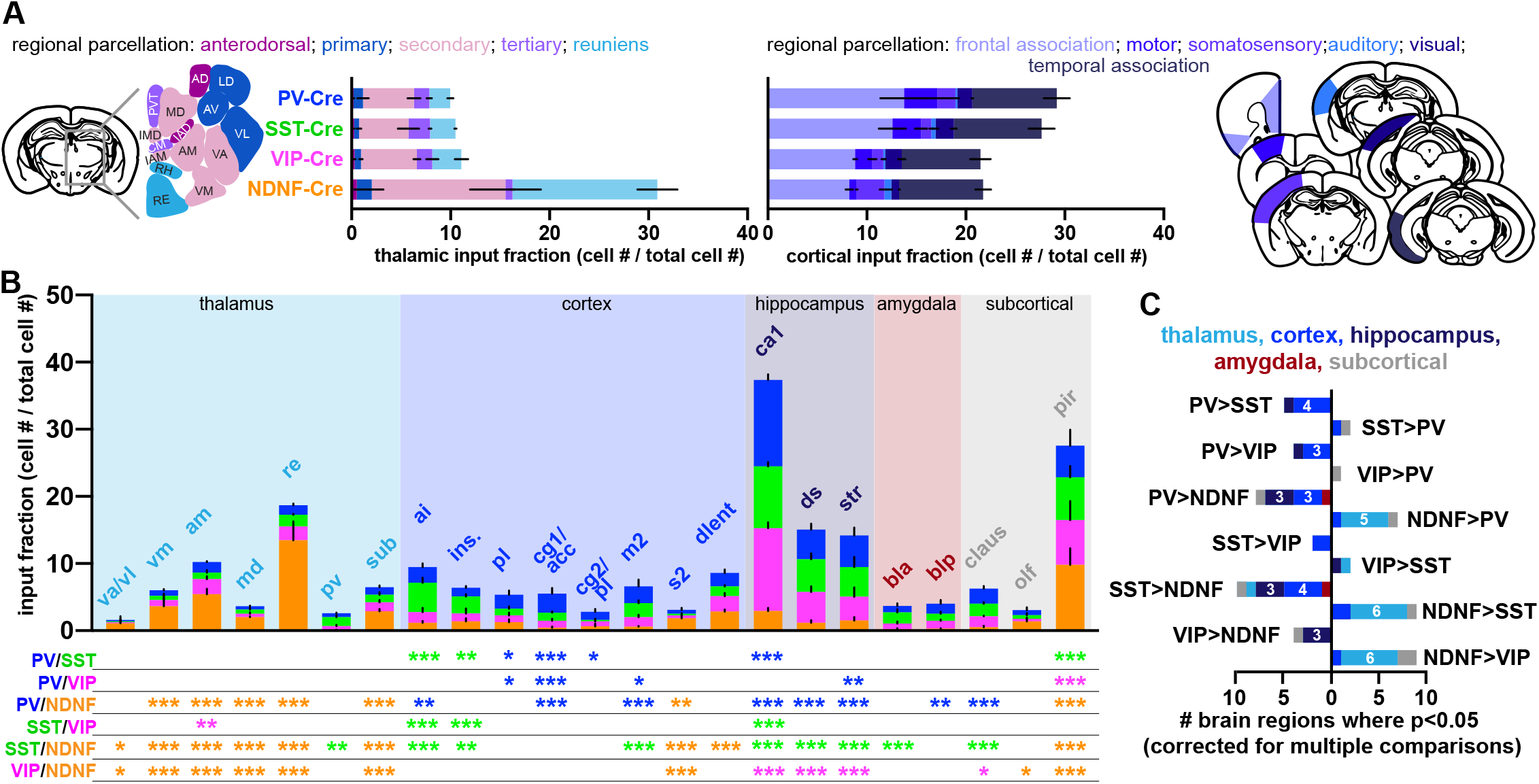
Differences in the input fraction among regions targeting IL interneurons. **A)** Proportion of inputs targeting IL interneurons from distinct thalamic groups (left) and cortical information processing modalities (right). **B)** Proportion of inputs from individual brain regions; the asterisks below denote the statistical result (*, p<0.05; **, p<0.005; and ***, p<0.0005 after adjustments for multiple comparisons) and the colors refer to the Cre driver line with the higher input fraction. **C)** The total number of individual brain regions that differ significantly across each pairwise comparison. Refer to Supplemental Figure 5 and Table S4 for specific details on the statistical comparisons and Table 1 for the full names of abbreviated brain regions.

**Table 1.** Region names and abbreviations associated with the results shown in Figure 5 and Figure 6.

| **<u>Region name</u>** | **<u>Abbreviation</u>** | **<u>Assignment</u>** |
| --- | --- | --- |
| ventral anterior /ventrolateral thalamic nucleus | va/vl | thalamus |
| ventromedial thalamic nucleus | vm | thalamus |
| anteromedial thalamic nucleus | am | thalamus |
| mediodorsal thalamic nucleus | md | thalamus |
| mediodorsal thalamic nucleus, lateral part | mdl | thalamus |
| reuniens thalamic nucleus | re | thalamus |
| paraventricular thalamic nucleus | pv | thalamus |
| submedius thalamic nucleus | sub | thalamus |
| lateral orbital frontal cortex | lo | cortex |
| anterior insular cortex | ai | cortex |
| insular cortex | ins | cortex |
| prelimbic frontal cortex | pl | cortex |
| infralimbic frontal cortex | il | cortex |
| cingulate cortex, area 1/ anterior cingulate cortex | cg1/acc | cortex |
| cingulate cortex, area 2/ prelimbic frontal cortex | cg2/pl | cortex |
| secondary somatosensory cortex | s2 | cortex |
| perirhinal cortex | prh | cortex |
| dorsolateral entorhinal cortex | dlent | cortex |
| field CA1 of the hippocampus | ca1 | hippocampus |
| dorsal subiculum | ds | hippocampus |
| ventral subiculum | vs | hippocampus |
| subiculum, transition area | str | hippocampus |
| basolateral amygdaloid nucleus, anterior part | bla | amygdala |
| basolateral amygdaloid nucleus, posterior part | blp | amygdala |
| claustrum | claus | subcortical |
| olfactory nerve layer | olf | subcortical |
| piriform cortex | pir | subcortical |
| dorsal peduncular cortex | dp | subcortical |
| lateral septal nucleus | ls | subcortical |

We next examined the proportion of input neurons across all brain regions (n=212). We found only a small number of regions that differed significantly different among PV-Cre, SST-Cre, and VIP-Cre neurons (PV vs. SST, 7 regions; PV vs. VIP, 5 regions; and SST vs. VIP, 4 regions). However, PV-Cre, SST-Cre, and VIP-Cre driver lines each had a greater number of regions that differed in comparison to NDNF-Cre (PV-Cre vs. NDNF, 15 regions; SST-Cre vs. NDNF, 18 regions; and VIP-Cre vs. NDNF, 12 regions) (**Figure 5B**). The largest individual input fraction to PV-Cre, SST-Cre, and VIP-Cre driver lines was from hippocampal area CA1 (~12%); however, the CA1 input fraction to NDNF-Cre neurons was only ~3% (p<0.0001 for each line vs. NDNF-Cre) (**Figure 5B**). In contrast, the largest input fraction to NDNF-Cre neurons was from RE (~13%), which was significantly greater than the RE input fraction to PV-Cre, SST-Cre, or VIP-Cre (~2%; p<0.0001 for each vs. NDNF). Moreover, we found a systematic underrepresentation of midline thalamic neurons (e.g., RE, AM, Subthalamic, VM, Md-Th) targeting PV-Cre, SST-Cre, and VIP-Cre driver lines relative to NDNF-Cre, and a corresponding overrepresentation of hippocampal-associated regions (e.g., CA1, dorsal subiculum, and subicular transition region) targeting PV-Cre, SST-Cre, and VIP-Cre relative to NDNF-Cre (**Figure 5C**). Consistent with this notion, a principal component analysis revealed hippocampal and thalamic regions were among the top regions that contributed variability to the interneuron Cre driver line datasets (**Supplementary Figure 5**).

We corroborated these differences among the Cre driver lines by using an orthogonal, correlation-based approach. In this approach, we restricted our analysis to regions in which at least one Cre driver line received > 1% of the input fraction; this enabled us to avoid spuriously high correlations across the lines on account of the large numbers of afferent regions with near-zero values. We computed pairwise correlations (two-tailed Spearman’s r) between Cre driver lines and compared these correlations to simulated values from shuffled datasets (**Supplementary Figure 5**). The correlations among PV-Cre, SST-Cre, and VIP-Cre driver lines were high (approximately r= 0.75 for each line) and within the shuffled distributions; however, the correlations between each of these driver lines compared to NDNF-Cre was substantially weaker than expected (i.e., outside the expected distribution of the shuffled datasets; PV vs. NDNF: Spearman’s r= 0.22; SST vs. NDNF: Spearman’s r= 0.19; VIP vs. NDNF: Spearman’s r= 0.35) (**Supplementary Figure 5**). Collectively, these results reveal a dramatic shift between the primary afferent regions targeting IL cortical PV/SST/VIP interneurons from those targeting IL NDNF interneurons.

### Differences among afferent regions targeting IL excitatory projection neurons

We performed similar analyses to determine the differential targeting of IL afferents to excitatory projection neurons. As in the interneuron datasets, we found that the largest proportion of input from the thalamus belonged to regions in the secondary thalamic group, and that the proportion of secondary thalamic input differed across projection classes. Specifically, the proportion of secondary thalamic input targeting LEC- or PAG-projecting neurons was approximately ~two-fold greater than the than that targeting to BLA-projecting neurons and five-fold greater than those targeting RE-projecting neurons (**Figure 6A**). Unlike the large differences in input fraction from RE evident in the interneuron dataset, the proportion of input from RE to IL projection classes was similar. Within cortical information processing modalities that project to IL, LEC had larger input fractions from frontal cortical areas relative to both BLA and RE-projecting neurons; in contrast, BLA and RE-projecting neurons had significantly greater input fractions from temporal association cortices than LEC-projecting neurons (**Figure 6A**).

**Figure 6.**
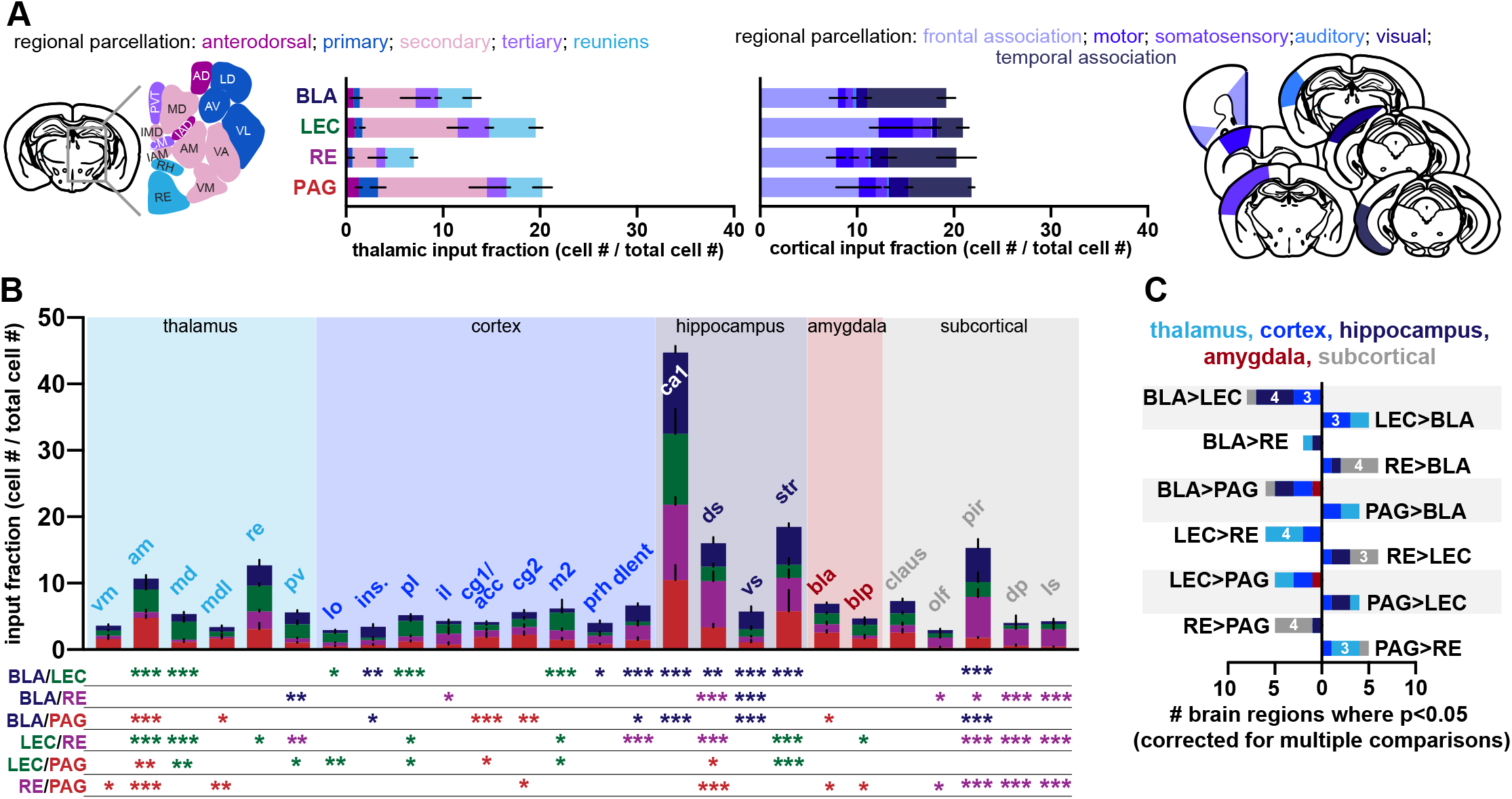
Differences in the input fraction among regions targeting IL projection neurons. **A)** Proportion of inputs targeting IL projection neurons from distinct thalamic groups (left) and cortical information processing modalities (right). **B)** Proportion of inputs from individual brain regions; the asterisks below denote the statistical result (*, p<0.05; **, p<0.005; and ***, p<0.0005 after adjustments for multiple comparisons) and the colors refer to the projection with the higher input fraction. **C)** The total number of individual brain regions that differ significantly across each comparison. Refer to Supplemental Table S5 for specific details on the statistical comparisons and Table 1 for the full names of abbreviated brain regions.

We next examined the input fractions from all brain regions to excitatory projection neurons. Across all presynaptic regions (n=212), the number of regions that differed in input fraction among the projection classes was similar to the interneuron datasets (BLA vs. LEC, 13 regions; BLA vs RE, 8 regions; BLA vs PAG, 10 regions; LEC vs RE, 13 regions, LEC vs PAG, 8 regions; RE vs PAG, 11 regions). The regions in which input fractions differed significantly among the projection classes largely overlapped with those identified in the interneuron datasets across the thalamus (e.g., RE, AM, PV, Md-Th), cortex (e.g., PL, M2 and DLENT), and hippocampus (e.g., CA1, dorsal subiculum, and subicular transition region) (**Figure 6B**). Although the effect sizes were more moderate than those found across the IL interneuron classes, systematic differences in the proportion of inputs from thalamus and hippocampus to projection neurons were evident. For example, BLA and LEC projecting neurons differed significantly from each other across 13 brain locations including hippocampus (4 regions) and thalamus (2 regions); LEC-projecting neurons had a greater input fraction from both thalamic regions (AM and Md-Th), while BLA projecting neurons had a greater input fraction from each of the 4 hippocampal regions (CA1, dorsal subiculum, ventral subiculum, and subicular transition region). Similarly, LEC- and RE-projecting neurons differed from each other in 12 brain regions, including hippocampus (2 regions) and thalamus (4 regions); LEC had greater input from all four thalamic regions (AM, Md-Th, PV, and RE) while RE projecting neurons had greater input from the CA1 and dorsal subiculum in the hippocampus. In contrast, BLA and RE projecting neurons had the fewest differences and did not show a systematic over- or underrepresentation of thalamic or hippocampal inputs (**Figure 6B,C**).

In support of differential connectivity between thalamic and hippocampal regions to IL projection classes, a PCA identified CA1 and AM as the top regions that contribute to the variance among the datasets (28% and 17% of the total variance in PC1, respectively) (**Supplementary Figure 6**). The same correlation-based approach that was used to examine the interneuron classes revealed that LEC and RE-projecting neurons had correlations lower than would be expected by chance (Spearman’s r= 0.19), while differences between PAG and RE-projection neurons matched the lower 2.5% cutoff we used as a bound for statistical significance (**Supplementary Figure 6**).

Many of the same regional differences that were evident among the interneuron classes or among the projection neuron classes remained when we analyzed all eight cell classes at once, demonstrating their statistical robustness (**Supplementary Figure 7**). Collectively, our results provide strong evidence that both inhibitory and excitatory projection neurons in IL receive input from a widespread and overlapping set of afferent brain regions; yet the proportion of afferent neurons that target these distinct postsynaptic neurons differ, with prominent differences most evident between hippocampal and thalamic regions. Interestingly, the wiring patterns appear to provide a structural basis for two pathway-specific IL microcircuits (schematized in **Figure 7A,B**).

**Figure 7.**
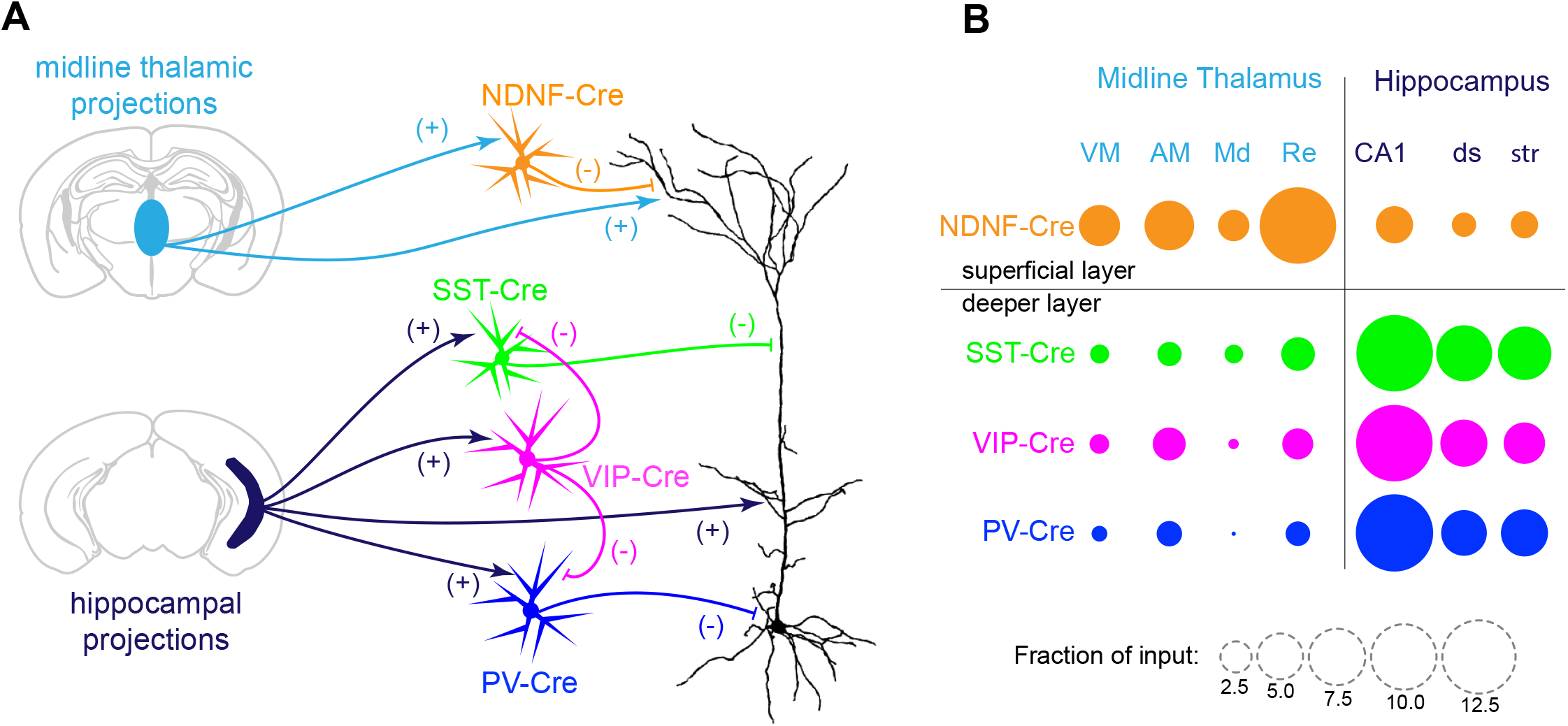
Biased cellular targeting of thalamic and hippocamapl afferents suggest the presence of two prominent IL microcircuits. **A)** The spatial targeting of thalamic or hippocampal excitatory synapses (+) onto the distal or proximal dendrites of pyramidal cells, coupled with the spatial targeting of inhibitory synapses (-) from these interneuron classes, suggests two prominent feedforward microcircuits controlling the excitability of IL projection neurons. **B)** A graphical depiction of the differential targeting of these afferents to IL interneurons.

## Discussion

Exactly how networks in the frontal cortex signal task-relevant features to support optimal decision making remains unknown. The frontal cortex integrates information from a widespread number of sensory, motor, emotive, and memory-related brain regions to support the flexible selection of goal-directed behaviors. Previous work has defined the regions that project to or receive projections from rodent frontal cortex (Gabbott et al., 2005; Hoover and Vertes, 2007), though our knowledge of the cell-type-specific wiring patterns remains incomplete. Here, we used monosynaptic, retrograde rabies tracing to map the afferent pathways that converge onto specific frontal cortical inhibitory and excitatory cell classes. This paper reports the analyses of these datasets and is accompanied by a supporting website (http://raisin.janelia.org, which we will continue to update during the peer review process) where visitors can make comparisons between inputs to postsynaptic cell classes and download raw data files.

The IL interneurons we examined here all appear to receive the largest fraction of their inputs from long-range afferent circuits. This result differs markedly from recently published work using similar rabies strategies to map connectivity onto PV-Cre, SST-Cre, and VIP-Cre interneurons (Ahrlund-Richter et al., 2019; Sun et al., 2019), which reported that the overwhelming fraction of neurons targeting PV-Cre, SST-Cre, and VIP-Cre interneurons were made by local frontal cortical neurons. However, both experiments used a different strain of modified rabies, SAD-B19ΔG, which has been found to label afferent neurons with a lower efficiency than the CVS-N2cΔG strain used here (see (Reardon et al., 2016). Thus, our results from both interneurons and projection neurons produce anatomical maps that look substantially different from the existing rabies maps and are more consistent with results provided by traditional retrograde tracers and channelrhodopsin-assisted mapping (CRACM) (Anastasiades et al., 2020; Liu and Carter, 2018).

Using Cre driver lines, we find that PV-Cre, SST-Cre, and VIP-Cre interneurons shared highly correlated patterns of afferent inputs across the brain, with the predominant fraction of presynaptic neurons originating in hippocampal region CA1 and the subiculum. Despite such broad similarity in their input patterns, the strength of these synaptic connections (Liu et al., 2020), the unique intrinsic biophysical properties of the postsynaptic neuron classes (e.g., fast-spiking vs adapting, etc.) and the specificity of their subsequent postsynaptic targets (e.g., forming synapses onto pyramidal cells or interneurons, or to somatic or dendritic compartments, etc.) (reviewed in (Tremblay et al., 2016) permit these different classes to produce different forms of inhibition within the IL microcircuit. NDNF-Cre interneurons, which reside exclusively in layer 1, had input patterns that were distinctly different than PV-Cre, SST-Cre, and VIP-Cre interneurons. The primary driver of these neurons appeared to be RE rather than CA1, and the correlations between afferent patterns targeting NDNF and the other interneuron classes were low. Collectively, these results provide strong evidence that NDNF interneurons provide a unique form of feedforward inhibition in IL.

We created brain-wide maps of afferent input onto excitatory projection neurons using a RetroAAV approach (Tervo et al., 2016) and similar to (Schwarz et al., 2015). Each of the IL projection neurons we examined received their greatest input fraction from area CA1 of the hippocampus, which appears consistent with channelrhodopsin-assisted mapping (CRACM) of CA1 connections onto IL projection neurons (Liu and Carter, 2018) and with the extremely high density of CA1 axons localized to the deeper layers of IL (Liu and Carter, 2018). Despite the large input fraction from CA1, a hippocampus-thalamus input dichotomy remained evident among the projection classes. For example, compared to RE-projecting neurons, LEC-projecting IL neurons receive a greater fraction of inputs from multiple thalamic regions and fewer inputs from hippocampus. This pattern (i.e., fewer from hippocampus; more from thalamus and *vice versa*) also held across more stringent comparisons between projection neuron pairs sharing the same somatic layer and dendritic morphology (e.g., LEC-projecting neurons vs. BLA-projecting neurons). Thus, our data suggest that variation in the functional outputs of IL projection neurons may be driven by the different proportional connectivity with hippocampal and thalamic afferent circuitry.

The cellular targeting of midline thalamic regions to layer 1 NDNF-Cre interneurons, and the corresponding spatial targeting of these axons to the most distal portions of the dendritic arbor of pyramidal cells (Anastasiades et al., 2020), creates a local microcircuit in which feedforward inhibition and feedforward excitation interact directly within the distal apical tuft of pyramidal cells. This motif is consistent with CRACM studies in which mouse frontal cortical NDNF interneurons receive strong and robust input from midline thalamus and control distal dendritic Ca^2+^ electrogenesis in frontal cortical pyramidal cells (Anastasiades et al., 2020). The cellular targeting of hippocampal output (e.g., from CA1) to the deeper layers of IL along with interneurons that target the perisomatic (i.e., from PV-Cre neurons) and apical branches (i.e., from SST-Cre neurons) of the pyramidal cell arbor creates a second ‘push-pull’ microcircuit in the deeper cortical layers. VIP interneurons, known to preferentially inhibit SST interneurons (Pfeffer et al., 2013), may be driven by CA1 to provide a tunable form of inhibition within this hippocampal-to-IL microcircuit. Determining how these microcircuits interact to shape action potential output, plasticity, and feature selectivity will be a major advance toward understanding the functional role of cell types in frontal cortical-dependent behaviors.

Compared to CRACM, which typically tests a single pathway, our rabies maps provide a cell-type specific wiring diagram of all connected presynaptic partners across the brain (i.e., a “cell-type-specific projectome”). However, the number of synaptic connections made by single axonal branches, single neurons, or whole pathways onto postsynaptic targets remains unknown, as does their spatial organization on the dendrites, including branch-specific targeting or intrabranch clustered or distributed patterns. Despite these shortcomings, our results are the first to suggest a systematic hippocampal-thalamic dichotomy that characterizes connectivity onto spatially intermingled neuronal classes in the mouse ventromedial frontal cortex. The patterns of connectivity we find here differ from those that appear to characterize connectivity in the hippocampus, where inputs to CA1 pyramidal cells from CA3, LEC, or MEC appear to depend on the spatial position of the postsynaptic neuron in area CA1 (i.e., connectivity gradients map well onto spatial gradients) rather than its projection target *per se*.

Future experiments should seek to identify specific functions for these IL afferent pathways to determine how their activity influences information transforms within frontal cortical circuitry. Given the large divergence of postsynaptic targets from even a single pathway, these results may spur new approaches to manipulate specific afferent connections rather than whole pathways *in toto*. The emergence of such technology will provide new ways to dissect how the structural organization of synaptic connectivity within cortical circuits contributes to their emergent functions.

## Methods

### Mice

All experiments were conducted in accordance with NIH guidelines and with approval of the Janelia Institutional Animal Care and Use Committee (Protocol 14-118). Adult C57BL/6J male mice (between 10-16 weeks of age) were used for the tracer and AAV experiments in Figure S1, for the overlap of projection neurons in Figure 2 and S2, for the CVS-N2cΔG EGFP(EnvA) control experiments designed to test the leak of AAV and RabV reagents in Figure S3, and for all experiments that used AAVRetro-Cre to map afferent neurons targeting IL projection neurons. Homozygous Rosa26-LSL_GFP-H2B mice were used for the AAVRetro-Cre experiments shown in Figure S1. Heterozygous transgenic mice with an ires-Cre coding sequence inserted into the promoter region of the *pv* (Hippenmeyer et al., 2005), *sst* (Royer et al., 2012), *vip* (Taniguchi et al., 2011), or *ndnf* (Tasic et al., 2016) loci were used to gain access to distinct sets of interneurons. In all experiments, mice were single housed after surgery in a 12-h/12 h light/dark cycle with food and water available ad libitum. Mice were assigned to experimental conditions based upon their availability.

### In situ hybridization, dual recombinase-dependent reporter labeling, and confocal microscopy

To examine the specificity, efficiency, and cellular overlap of each transgenic Cre driver line in IL (shown in Figure 1), 20 μm-thick cryostat sections of perfusion fixed brains were used for RNAscope fluorescent triple-label *in situ* hybridization according to the manufacturer’s instructions and with commercially-available reagents (Advanced Cell Diagnostics; similar to (Bloss et al., 2016)). Tiled z-stacks of IL were acquired using a Zeiss 710 confocal microscope equipped with a 20x objective and ZEN software, and images were quantified using the cell counter plugin for Fiji. To measure the cellular overlap of IL neurons projecting to different target sites, 150 μm-thick sections containing IL tdTomato or GFP-expressing projection neurons were imaged using a Zeiss 710 confocal microscope equipped with a 40x objective and ZEN software. The fraction of single and dual labeled neurons was counted using the cell counter plugin for Fiji. In both experiments, the distance of the labeled cell from the pial surface was recorded to obtain the data in Figure S2. For the dendritic morphological reconstructions, interneurons and projection neurons labeled by Cre- or FLPo-dependent reporter viruses were acquired with a Zeiss 710 confocal microscope equipped with a 40x objective and ZEN software, TIFF stacks were imported into NeuronStudio, and dendritic morphologies were manually reconstructed and analyzed using a Sholl analysis.

### Viruses and Intracranial Injections

Viruses were prepared at Janelia Research Campus, and titers were as follows: AAVRetro-Cre, 1e13 GC/mL; AAVRetro-FLPo, 1e13 GC/mL; AAVRetro-tdTomato-H2B, 1e13 GC/mL; AAV2/1-CamKII-Cre, 4e13 GC/mL; AAV2/1-CAG-flex-*rev*-GFP, 1e13 GC/mL; AAV2/1-CAG-flex-*rev*-mKate2-T2A-N2c-G, 2e13 GC/mL; AAV2/1-CAG-flex-*rev*-mKate2-T2A-TVA, 1e13 GC/mL; and CVS-N2cΔG EGFP(EnvA), 1e9 IU/mL. Injection volumes were chosen to minimize spread into the adjacent motor or prelimbic cortices and were the following: WGA-555 (Sigma) and CTB-555, both 36 nL per site; AAVRetroCre, 27 nl per site. For IL injection of AAV2/1-CAG-flex-*rev*-mKate2-T2A-N2c-G and AAV2/1-CAG-flex-*rev*-mKate2-T2A-TVA, the two viral constructs were mixed at a 2:1 ratio. The methodology and details for intracranial injections of tracers and viruses was identical to that described in (Bloss et al., 2016). For experiments in which IL was targeted by a pair of injections (e.g., in all rabies experiments where an initial injection of AAVs was followed by injection of CVS-N2cΔG), the first IL injections of AAVs were made with the mouse at a 0° tilt, and the subsequent RabV was injected with the mouse at a 15° tilt to avoid inadvertent transduction of neurons along the first pipette tract. At all sites, high-titer viral suspension (18-54 nl) was injected over 5 minutes at the following coordinates (in mm relative to bregma, lateral relative to midline, and ventral relative to pial surface):

IL: (+1.75, 0.3, 2.25);

IL at 15° tilt: (+1.75, 0.95, 2.6);

BLA: (−1.6, 3.3, 4.1);

PAG, (−4.75, 0.5, 2 and 1.5);

LEC: (−4.2, 4.5, 2.5);

RE at 15° tilt (−1.2 and −1.5,1.2, 4.2).

### Tissue Processing and Imaging

Mice were sacrificed by transcardial perfusion at the following timepoints after intracranial injection: WGA, 24 hours; CTB, 1 week; AAVRetro-Cre (for Fig. S1), 2 weeks; AAVRetro-tdTomato-H2B, 2 weeks; AAV-*rev*-flex-GFP, 2 weeks; RabV, one week. At sacrifice, mice were transcardially perfused with 5 mLs of ice-cold 1% depolymerized paraformaldehyde in 0.1M PB (pH 7.3), followed by 50 mls of ice-cold 4% paraformaldehyde in 0.1M PB (pH 7.3) at 10 mLs/min. Brains were postfixed in the same fixative overnight at 4°C, then transferred to 0.1M PB, cut into sequential 50 μm-thick vibratome sections, mounted onto microscope slides and coverslipped with Vectashield containing DAPI.

Images of coronal sections (from 2.68 mm anterior to Bregma to −4.6mm posterior to Bregma) were collected on a TissueFaxs 200 confocal microscope (TissueGnostics, Vienna, Austria) comprising an X-Light V2 spinning disk confocal imaging system (CrestOptics, Rome, Italy) built on an Axio Imager.Z2 microscope (Carl Zeiss Microscopy, White Plains, NY) and equipped with a 10x, 0.3 NA objective (Zeiss) and an Zyla 5.5 sCMOS camera (Andor, Belfast, UK). DAPI, EGFP, and mKate2 were excited with a Spectra X light engine (Lumencor, Beaverton, OR).

Z-stacks (3 μm step) were collected to collect data from the entire slice, but only maximum intensity projections were analyzed. Exposure times were kept constant throughout the study for all three channels, though the dynamic range of the exported 16-bit images was adjusted to correct for the mild differences in signal intensity between samples. Assembled slice montages were exported as 16-bit images then transformed to 8-bit images during alignment to (Paxinos and Franklin, 2004). Briefly, individual coronal slices were registered via affine transformations in trakEM2 (Fiji) using the tissue edges and various internal reference marks (e.g., ventricles or white-matter tracts). EGFP+ and mKate2+ neurons were counted and assigned to specific brain regions using intensity-based thresholds. False negative and Positive rates of this counting procedure are shown in Figure S3.

### Statistical Analyses

All statistical comparison and results are reported in Supplemental Tables S1-S8. Briefly, all two-way ANOVAs (e.g., in Fig 3D, Fig 4B-D, Fig 5A-B, Fig 6A-B, Fig S2,B Fig S2D, Fig S3C) used Tukey’s posthoc tests with adjustments for multiple comparisons, and all correlations (e.g., Fig 3E, Fig S3B, Fig S5C, and Fig S6C) used two-tailed nonparametric Spearman’s correlations; these datasets were teste for normality and analyzed using GraphPad Prism 8. Principal component analyses were run on R using the prcomp package. Figures in the paper depict mean ± SEM. The raw images and data for the figures is available at (http://dx.doi.org/10.6084/m9.figshare.13302731) and at (http://raisin.janelia.org).

## Author Contributions

EBB conceived the project and performed the experiments. KG and EBB analyzed the data. KG, NS, and EBB wrote the paper.

## Acknowledgements

We thank Drs. Carmen Robinett and Brett Mensh for critical comments on the manuscript; Dr. Kim Ritola for virus production; Sara Lindo and Sal Dilisio for surgical expertise; Monique Copeland for histology; and Deanna Otstot for mouse genotyping and breeding. We would like thank Jody Clements and Dave Davis Bennett for their work on the RAISIN website. This work was made possible by internal funding from the Howard Hughes Medical Institute and The Jackson Laboratory.

