## Supplemental Figures for "Hippocampal and thalamic afferents form distinct synaptic microcircuits in the mouse frontal cortex"

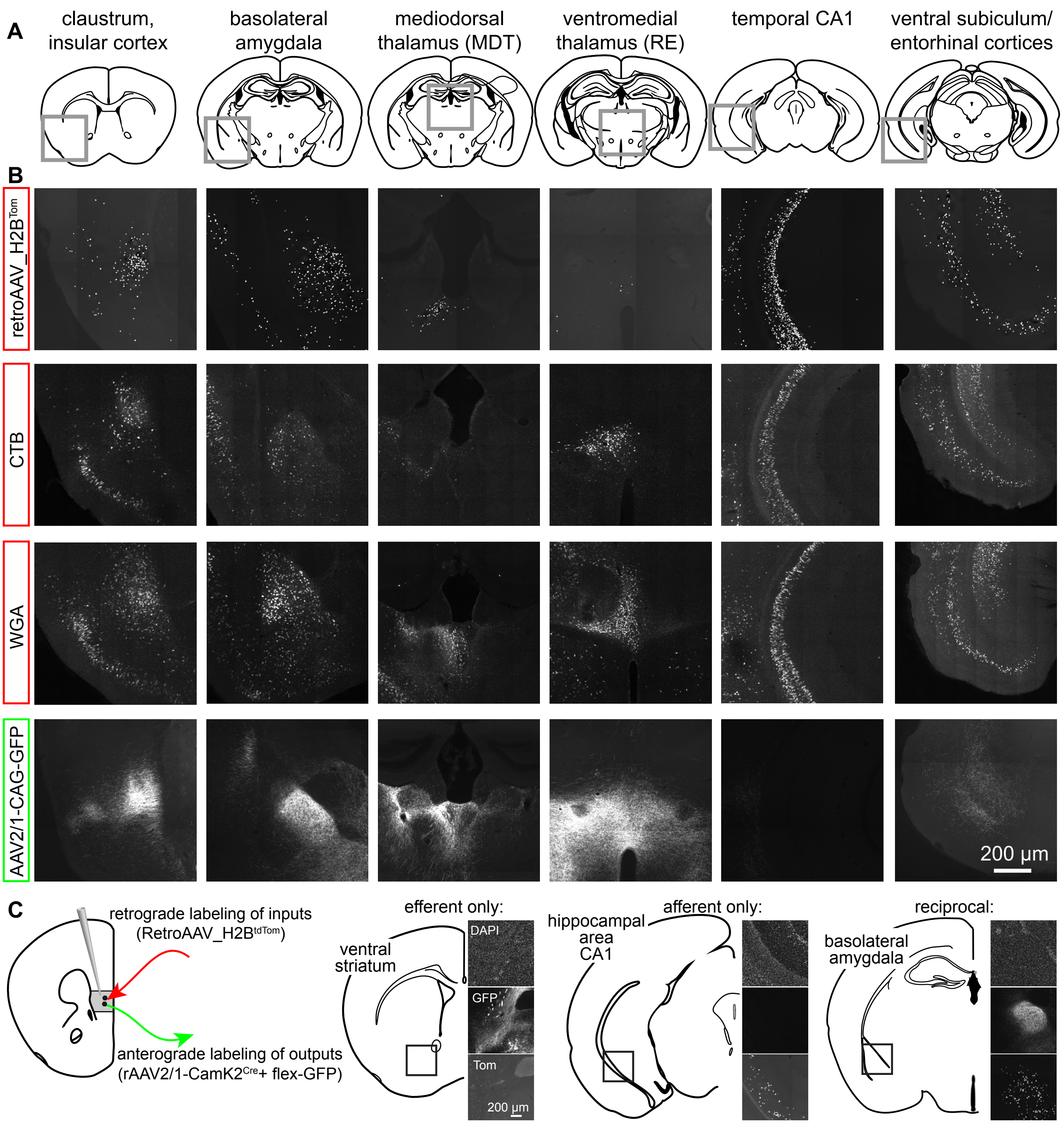


**Supplemental Figure 1. Maps of afferent and efferent pathways to mouse IL cortex.** A) Cortical and subcortical brain regions connected to IL. **B)** Images of the regions in A taken from experiments with IL injections of AAV-retro, CTB, WGA, and anterograde GFP (rows from top to bottom). **C)** Examples from dual labeling experiments that highlight pathways containing IL efferents only (e.g, ventral striatum), IL afferents only (e.g., area CA1), or pathways that are reciprocally connected (e.g., basolateral amygdala).

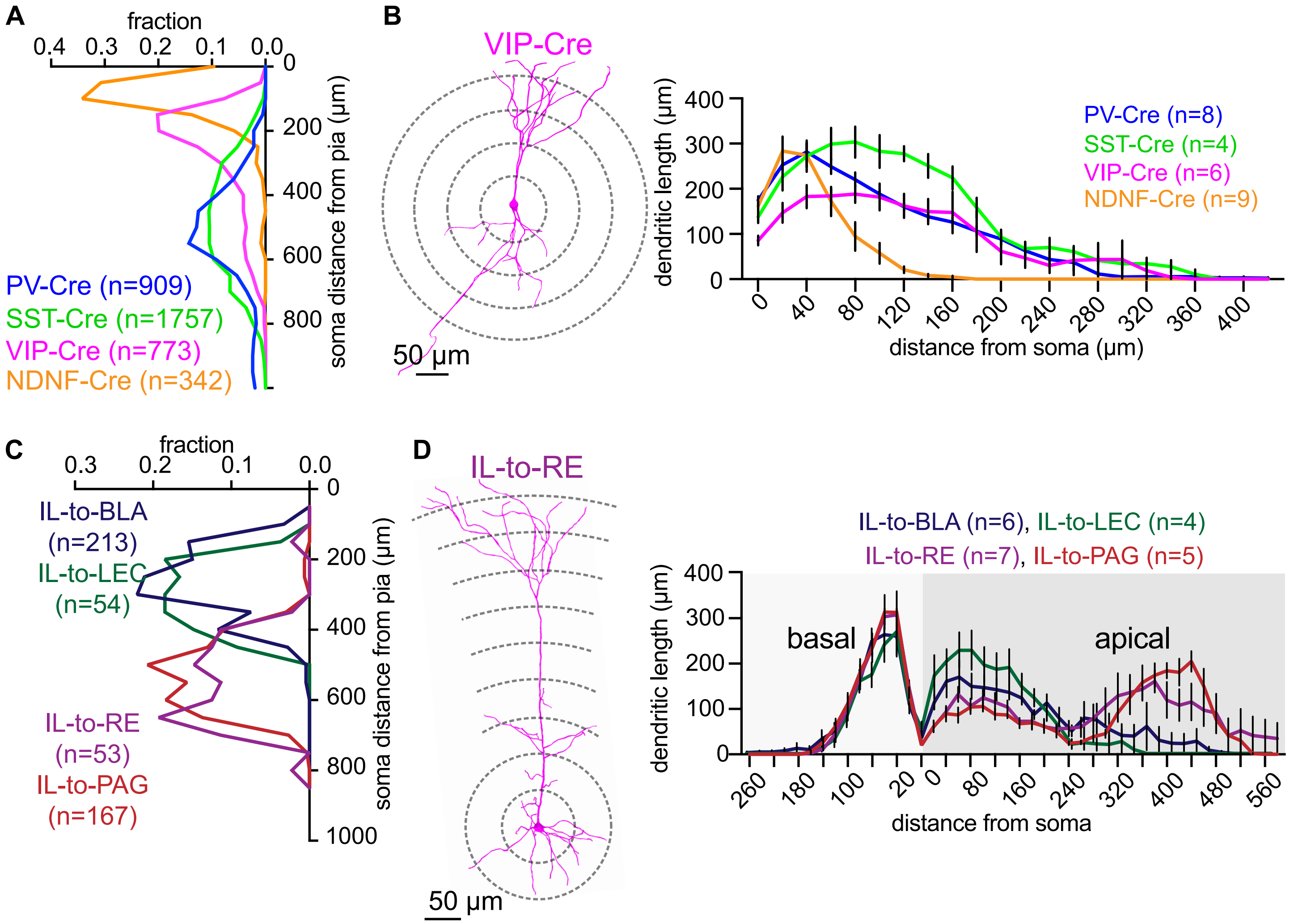


**Supplementary Figure 2. Somatic location and dendritic morphology of IL neurons. A)** Distance of the soma from the pial surface for each inhibitory interneuron Cre driver line (data taken from the *in situ* experiments). **B)** Sholl-style analysis of the morphology of their dendritic arbors. **C)** Somatic distance from the pial surface differed between superficial (i.e. BLA and LEC; dashed lines) and deep (i.e., RE and PAG; solid lines) projection neurons. **D)** Reconstruction of the dendritic morphologies of IL projection neurons. See Table S6 for the detailed results of statistical analyses.

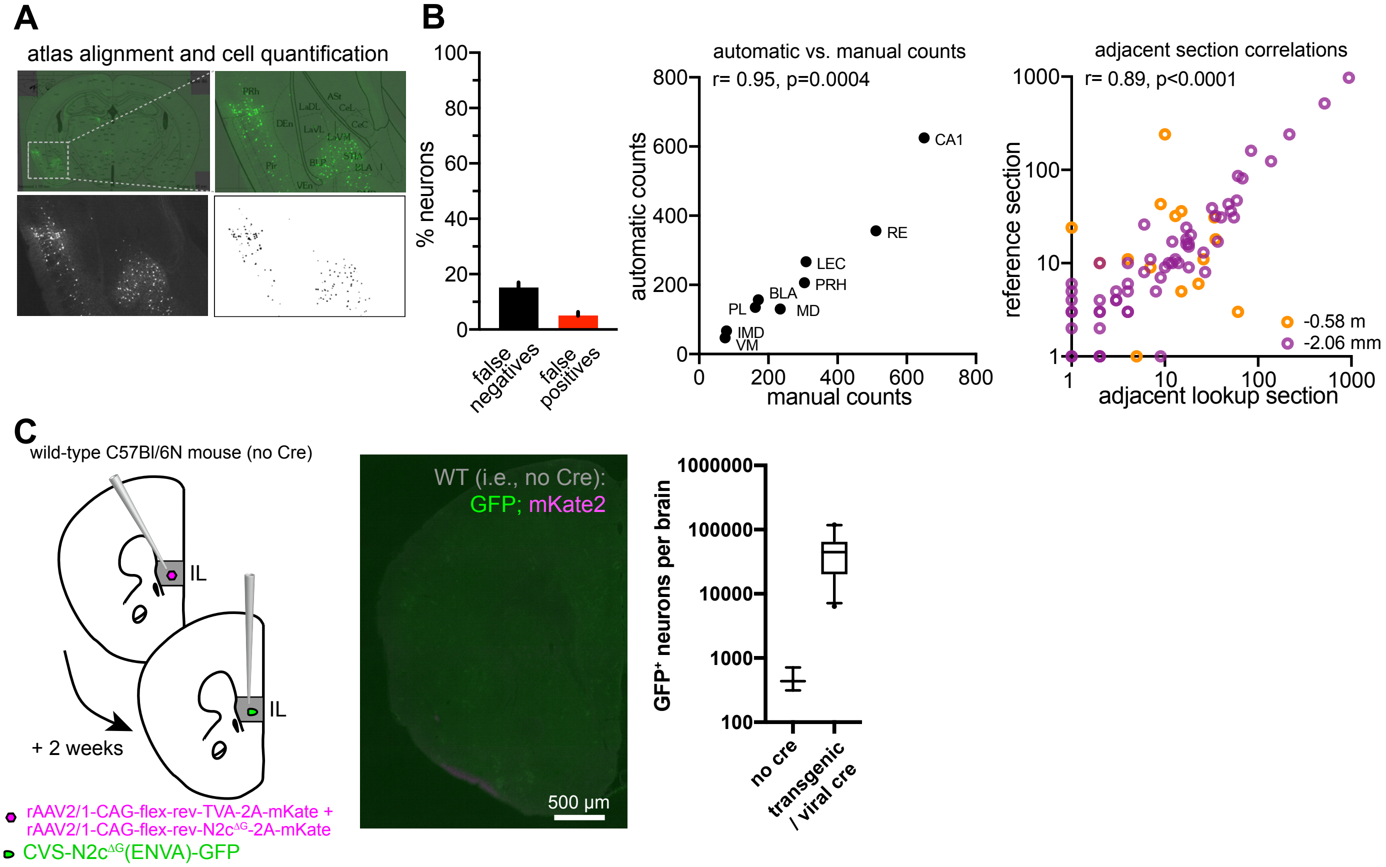


**Supplementary Figure 3. Control experiments for anatomical quantification of RabV-labeled EGFP⁺ neurons. A)** Coronal slices were imaged, aligned to reference sections from (Paxinos and Watson), and quantified using semi-automated cell counting algorithms in FIJI. **B)** False negative and false positive rates were determined by comparisons between manual and automated counts (left): strong correlations were found on the same section independent of regional cellular density, as well as between automated counts from neighboring sections assigned to the same coronal plane (two comparisons are shown, from -0.58 mm posterior from Bregma and from -2.06 mm posterior from Bregma). **C)** Control experiments were performed using mice (n=3) that did not express Cre to determine leak in the Cre-dependent reagents and RabV labeling, and these experiments revealed few EGFP⁺ neurons across the brains of these mice (~500 neurons/brain) relative to those from the experiments with transgenic or viral Cre expression (~50,000 neurons/brain). See Table S7 for the detailed results of statistical analyses.

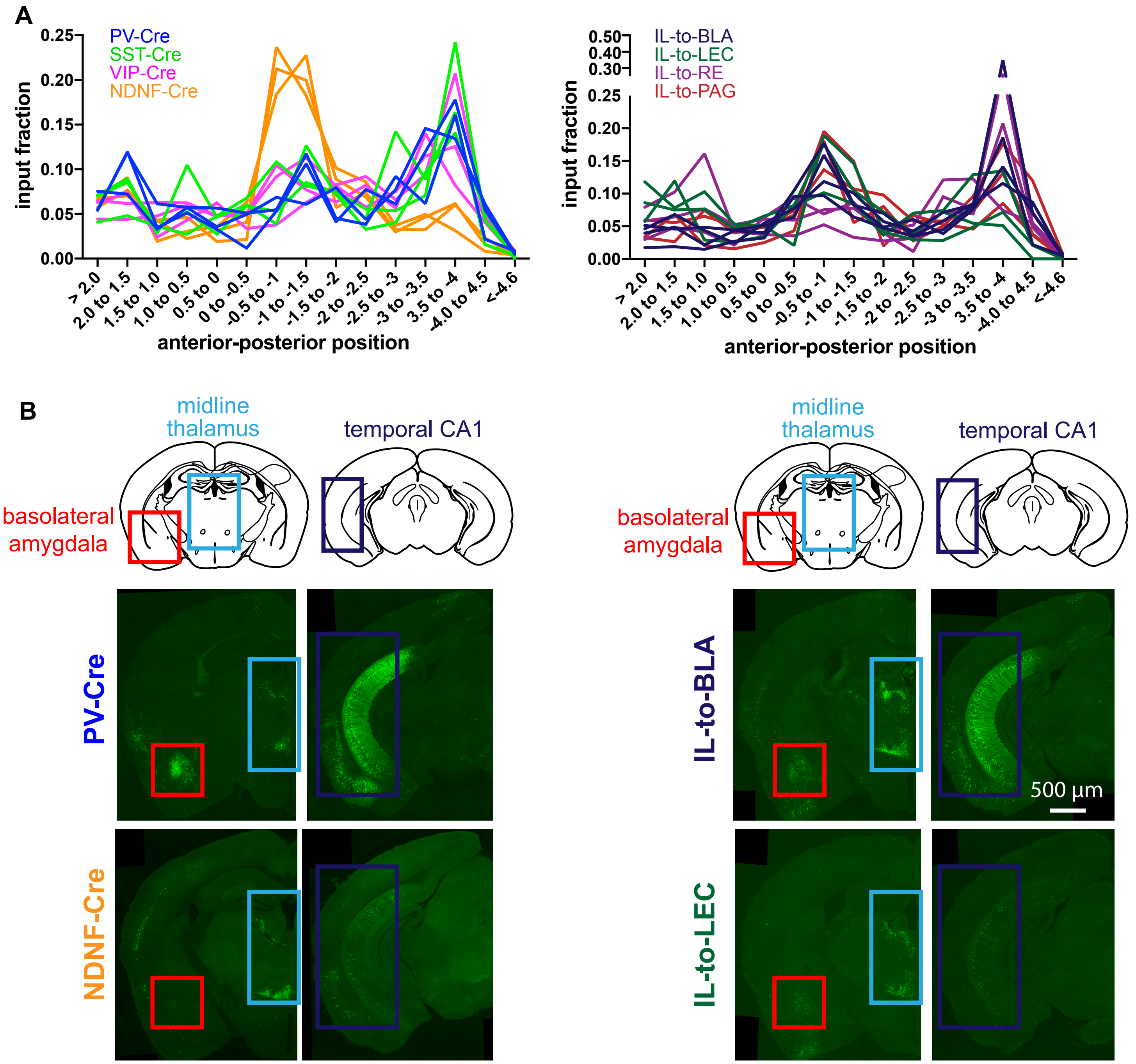


**Supplementary Figure 4. Spatial distribution of afferent neurons across individual replicate experiments.** A) Plots (as shown in Figure 4) with individual replicates shown. B, representative images from a PV-Cre and NDNF-Cre mouse (left) and IL-to-BLA and IL-to-LEC mouse (right) at the anterior peak and at the posterior peak.

A

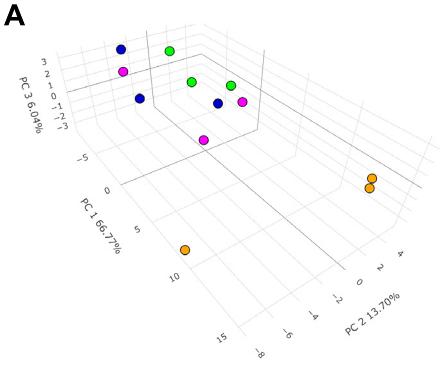


thalamus, cortex, hippocampus, amygdala, subcortical

| PC 1 | | PC 2 | | PC 3 | |
|---|---|---|---|---|---|
| RE | 44.3 | PIR | 59.2 | STR | 28.9 |
| CA1 | 24.0 | VM | 9.9 | DS | 12.6 |
| PIR | 11.1 | AM | 7.2 | RE | 12.1 |
| DS | 3.9 | PL | 3.0 | VM | 10.9 |
| AM | 3.5 | VA/VL | 2.7 | AI | 6.3 |
| STR | 3.0 | INS | 2.6 | PIR | 4.7 |
| VM | 1.5 | CA1 | 1.9 | CA1 | 4.6 |
| Subthal | 0.8 | AI | 1.8 | VA/VL | 2.9 |
| CG1/ACC | 0.8 | RE | 1.6 | AM | 2.9 |
| M2 | 0.7 | CG1/ACC | 1.5 | INS | 1.6 |
| Total Variance Explained (%) | 66.77 | | 13.7 | | 6.06 |

B

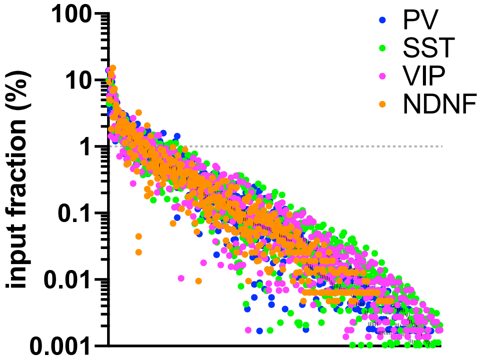


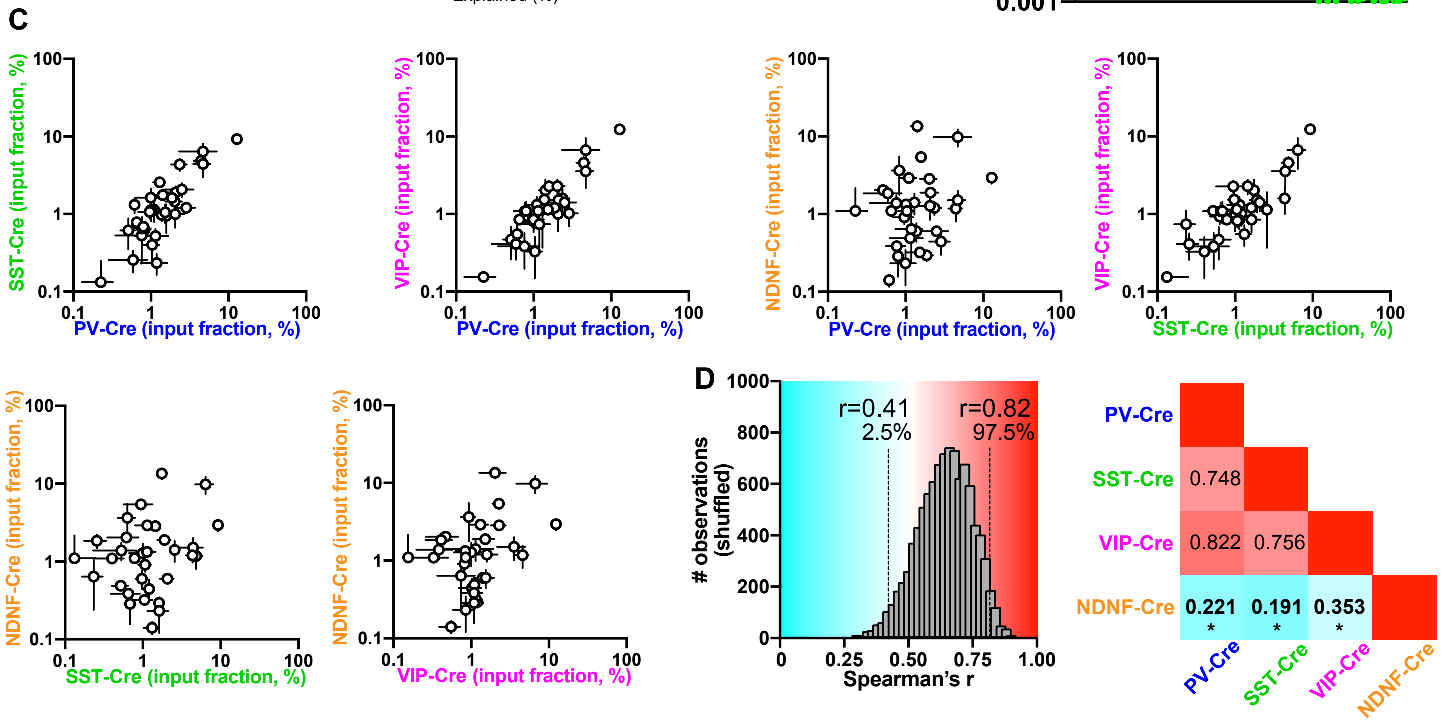


**Supplementary Figure 5. Differences in afferent regions targeting IL interneurons. A)** Principal component analysis on input fractions from afferent regions targeting IL interneurons, with individual replicates plotted in PCA space (left) and a table summarizing the top 10 regions for each of the first three components (right). **B)** Distribution of the input fraction from all 212 brain regions; the 1% threshold (dashed line) was used for the subsequent correlation analyses. **C)** Correlations across the Cre driver lines from the regions that meet the 1% threshold (n=32 regions). **D)** Spearman's r values from shuffled the datasets (n=10,000 shuffles) used to determine whether the observed correlations between pairs of Cre lines were higher or lower than expected (left, with the 2.5 and 97.5 percentiles that demarcate significance indicated by the dashed lines). A heat map of the actual correlations between each Cre driver line with the Spearman's r value within each color-coded box (right). See Table S8 for the detailed statistical results from these comparisons.

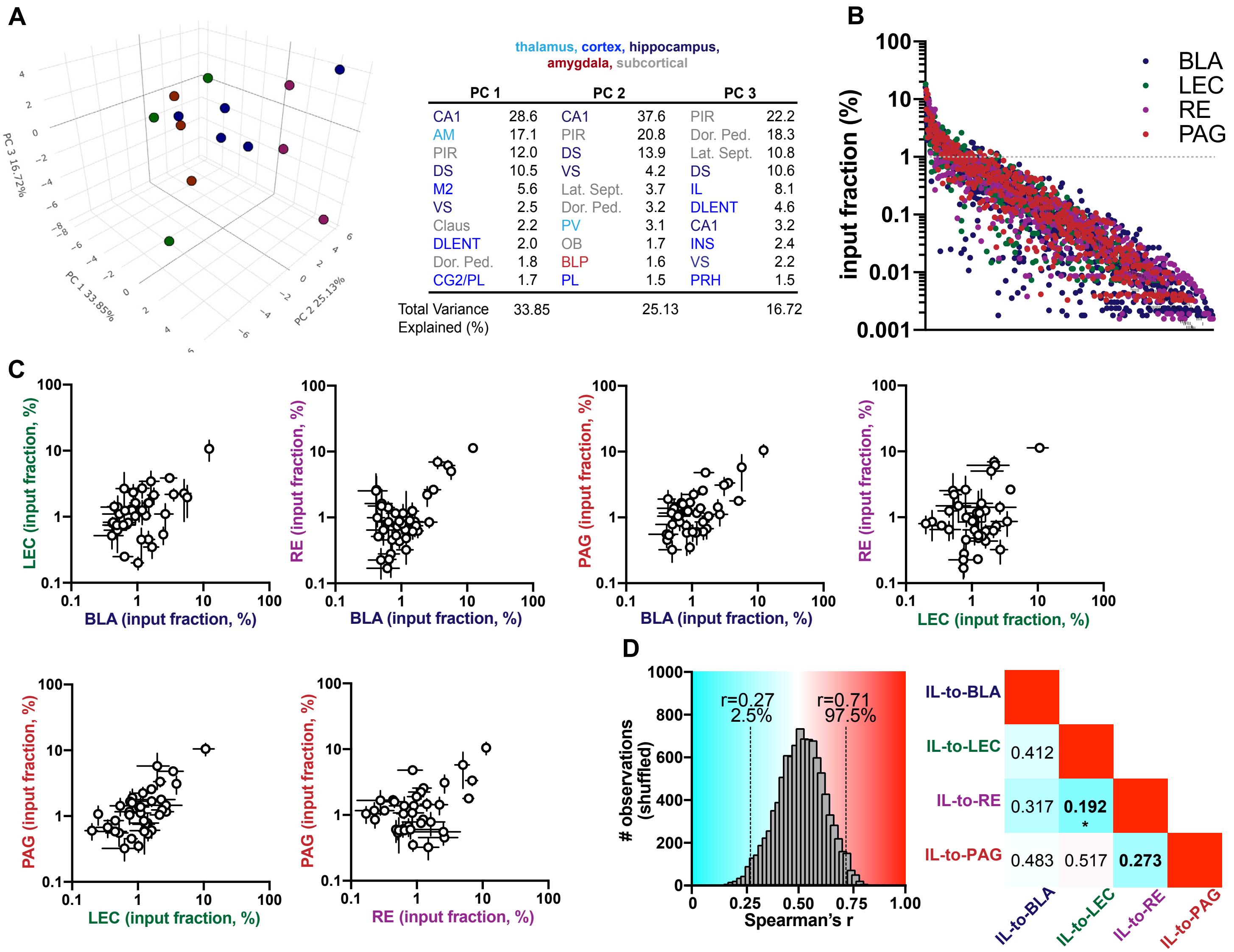


**Supplementary Figure 6. Differences in afferent regions targeting IL projection neurons. A)** Principal component analysis on input fractions from afferent regions targeting IL projection neurons, with individual replicates plotted in PCA space (left) and a table summarizing the top 10 regions for each of the first three components (right). **B)** Distribution of the input fraction from all 212 brain regions; the 1% threshold (dashed line) was used for the subsequent correlation analyses. **C)** Correlations across the projections neurons from the regions that meet the 1% threshold (n=38 regions). **D)** Spearman's r values from shuffled the datasets (n=10,000 shuffles) used to determine whether the observed correlations between pairs of projection classes were higher or lower than expected (left, with the 2.5 and 97.5 percentiles that demarcate significance indicated by the dashed lines). A heat map of the actual correlations between each projection class with the Spearman's r value within each color-coded box (right). See Table S9 for the detailed result of these statistical comparisons.

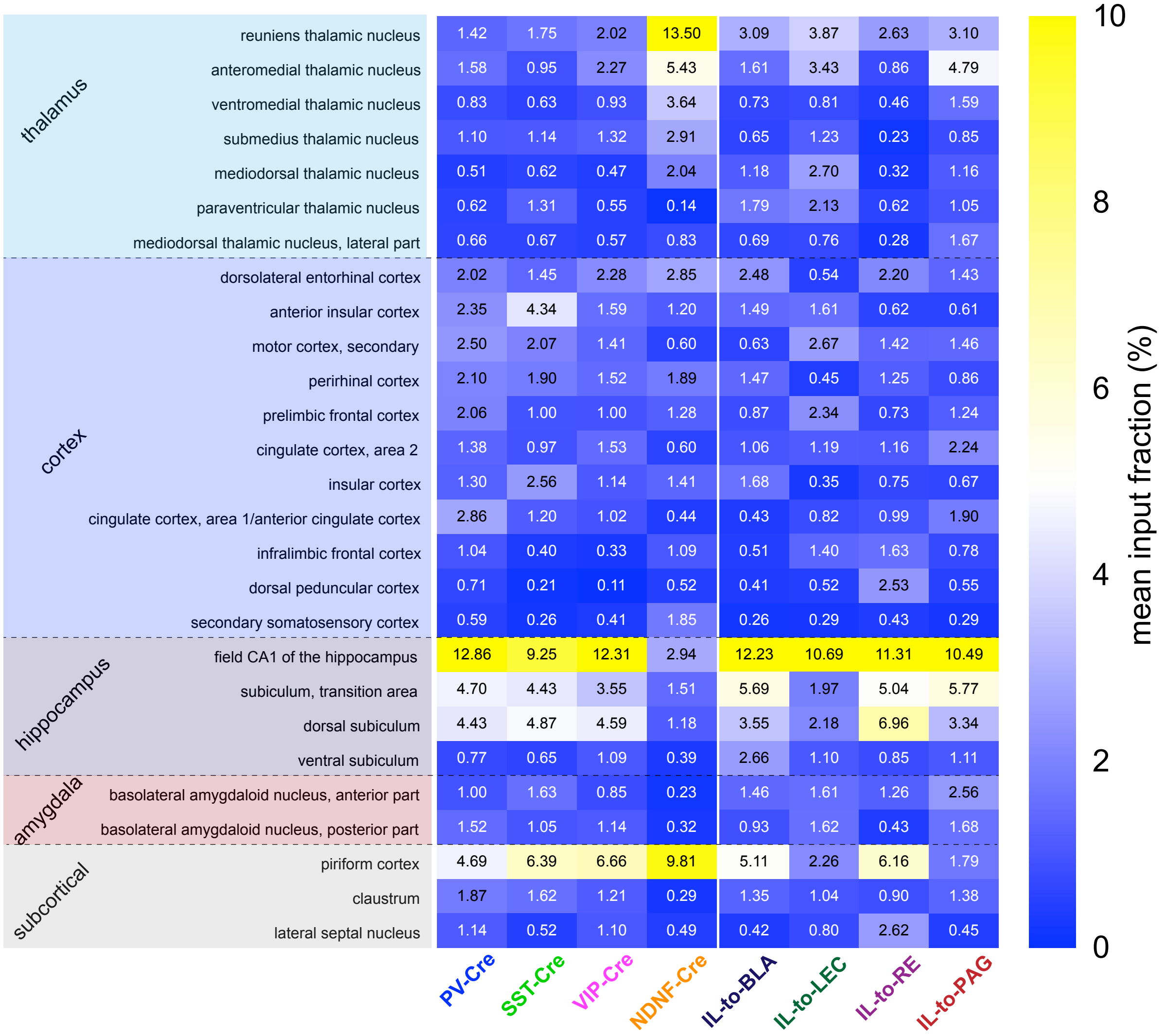


**Figure S7. Quantification of regional differences across all eight postsynaptic IL neuron classes.** A) Heat map showing the color-coded input fraction for 27 (of 212) regions which differed statistically across at least two postsynaptic cell classes after corrections for multiple comparisons. Regions are first organized by the parent brain structure (e.g., thalamus, cortex, etc) and then by average input fraction (from largest to smallest). The input fraction for each datapoint is indicated by text within the heatmap. See Table S10 for detailed statistical results from these comparisons.
