## Supplemental Tables for "Hippocampal and thalamic afferents form distinct synaptic microcircuits in the mouse frontal cortex"

**Table S1. A list of all regions analyzed in this study and the abbreviations used to refer to them in the figures.**

| Region name | Abbreviation |
|---|---|
| accumbens nucleus, core | accore |
| accumbens nucleus, shell | acbshell |
| amygdalohippocampal area, anterolateral part | ahial |
| amygdalohippocampal area, posteromedial part | ahipm |
| amygdalopiriform transition area | apir |
| anterior amygdaloid area | aa |
| anterior cortical amygdaloid area | aco |
| anterior hypothalamic area, anterior part | aha |
| anterior hypothalamic area, central part | ahc |
| anterior hypothalamic area, posterior part | ahp |
| anterior insular cortex | ai |
| anterior pretectal nucleus | apt |
| anterior pretectal nucleus, dorsal part | aptd |
| anterodorsal thalamic nucleus | ad |
| anterodorsal thalamic nucleus adjacent | ad2 |
| anteromedial thalamic nucleus | am |
| anteromedial thalamic nucleus, ventral part | amv |
| anteroventral thalamic nucleus | av |
| anteroventral thalamic nucleus, dorsomedial part | avdm |
| anteroventral thalamic nucleus, ventrolateral part | avvl |
| basolateral amygdaloid nucleus, anterior part | bla |
| basolateral amygdaloid nucleus, posterior part | blp |
| basolateral amygdaloid nucleus, ventral part | blv |
| basomedial amygdaloid nucleus | bma |
| caudate putamen (striatum) | cpu |
| caudomedial entorhinal cortex | cent |
| central amygdaloid nucleus, lateral division | cel |
| central amygdaloid nucleus, medial division | ceamy |
| central medial thalamic nucleus | cm |
| centrolateral thalamic nucleus | cl |
| cingulate cortex, area 1/anterior cingulate cortex | cg1/acc |
| cingulate cortex, area 2 | cg2 |
| cingulate cortex, area 2/ prelimbic frontal cortex | cg2/pl |
| claustrum | claus |
| cortex-amygdala transition zone | cxa |
| deep gray layer of the superior colliculus | dpg |
| deep white layer of the superior colliculus | dpwh |
| dorsal intermediate entorhinal cortex | dient |
| dorsal peduncular cortex | dp |

| | |
|---|---|
| dorsal raphe nucleus | dr |
| dorsal raphe nucleus - associated | dr ass |
| dorsal raphe nucleus, caudal part/dorsal raphe, interfascicular part | dr/cli |
| dorsal subiculum | ds |
| dorsolateral entorhinal cortex | dlent |
| dorsomedial hypothalamic nucleus | dmh |
| dorsomedial hypothalamic nucleus, compact part | dmc |
| ectorhinal cortex | ect |
| ethmoid thalamic nucleus | eth |
| extended amygdala- central part | eac |
| extended amygdala- medial part | eam |
| field CA1 of the hippocampus | ca1 |
| field CA2 of the hippocampus | ca2 |
| fields of Forel | ff |
| fornix | f |
| frontal association cortex | FrA |
| frontal cortex, area 3 | fr3 |
| globus pallidus | gp |
| infralimbic frontal cortex | il |
| insular cortex | ins |
| interanterodorsal thalamic nucleus | iad |
| interanteromedial thalamic nucleus | iam |
| intermediate gray layer of the superior colliculus | ing |
| intermediate white layer of the superior colliculus | inwh |
| intermediodorsal thalamic nucleus | imd |
| interstitial nucleus of Cajal, shell region | Insh/c |
| lateral amygdala | la |
| lateral amygdaloid nucleus, dorsolateral part | ladl |
| lateral amygdaloid nucleus, ventrolateral part | lavl |
| lateral amygdaloid nucleus, ventromedial part | lavm |
| lateral habenular nucleus | lhb |
| lateral hypothalamic area | lh |
| lateral hypothalamic area - adjacent | lhaa |
| lateral orbital frontal cortex | lo |
| lateral parietal association cortex | lpta |
| lateral posterior thalamic nucleus, laterorostral part | lplr |
| lateral posterior thalamic nucleus, mediorostral part | lpmr |
| lateral preoptic area | lpoa |
| lateral septal nucleus | ls |
| laterodorsal thalamic nucleus, dorsomedial part | lddm |
| laterodorsal thalamic nucleus, ventrolateral part | ldvl |
| lithoid nucleus | lth |
| magnocellular nucleus of the posterior commissure | mcpc |

| | |
|---|---|
| medial amygdaloid nucleus, anterior part | mea |
| medial amygdaloid nucleus, anterodorsal | mead |
| medial amygdaloid nucleus, posterodorsal part | mepd/v |
| medial entorhinal cortex | ment |
| medial habenular nucleus | mhb |
| medial orbital frontal cortex | mo |
| medial parietal association cortex | mpta |
| medial preoptic area | mpa |
| medial pretectal nucleus | mpt |
| medial septal nucleus | ms |
| medial terminal nucleus | mt |
| median raphe nucleus | mnr |
| mediodorsal thalamic nucleus | md |
| mediodorsal thalamic nucleus, central part | mdc |
| mediodorsal thalamic nucleus, lateral part | mdl |
| mediodorsal thalamic nucleus, medial part | mdm |
| mesencephalic reticular formation | mrt |
| motor cortex, primary | m1 |
| motor cortex, secondary | m2 |
| nucleus of Darkschewitsch | dk |
| nucleus of the horizontal limb of the diagonal band | hdb |
| nucleus of the optic | OT |
| nucleus of the posterior commissure | pcom |
| nucleus of the vertical limb of the diagonal band | vdb |
| nucleus of the vertical limb of the diagonal band/medial septum | vbd/ms |
| oculomotor nerve | 3n |
| oculomotor nucleus, parvicellular part | 3pc |
| olfactory nerve layer | olf |
| olivary pretectal nucleus | opt |
| optic nerve layer of the superior colliculus | op |
| oval paracentral thalamic nucleus | opc |
| p1 reticular formation | p1rt |
| parabrachial pigmented nucleus of the VTA | pbp |
| paracentral thalamic nucleus | pc |
| parafascicular thalamic nucleus | pf |
| paramedian raphe nucleus | pmnr |
| pararubral nucleus | par |
| parasubiculum | pas |
| paratenial thalamic nucleus | pt |
| paratrochlear nucleus | pa4 |
| paraventricular thalamic nucleus | pv |
| parietal cortex, posterior area, dorsal part | ptpd |
| parietal cortex, posterior area, rostral part | ptpr |
| peduncular part of lateral hypothalamus | plh |

| | |
|---|---|
| peduncular part of lateral hypothalamus/lateral hypothalamus | plh/lh |
| pedunculotegmental nucleus | ptg |
| periaqueductal gray | pag |
| perirhinal cortex | prh |
| piriform cortex | pir |
| pontine reticular nucleus, oral part | pno |
| posterior hypothalamic nucleus | ph |
| posterior pretectal nucleus | ppt |
| posterior thalamic nuclear group | po |
| posterolateral cortical amygdaloid area | plco |
| posteromedial cortical amygdaloid area | pmco |
| posteromedian thalamic nucleus | pomn |
| postsubiculum | post |
| precommissural nucleus | prc |
| precuneiform area | prcnf |
| prelimbic frontal cortex | pl |
| premammillary nucleus, dorsal part | pmd |
| premammillary nucleus, ventral part | pmv |
| prepositus nucleus | pr |
| presubiculum | prs |
| primary auditory cortex | au1 |
| primary somatosensory cortex | s1 |
| primary somatosensory cortex, barrel field | s1bf |
| primary somatosensory cortex, dysgranular zone | s1dz |
| primary somatosensory cortex, forelimb region | s1fl |
| primary somatosensory cortex, hindlimb region | s1hl |
| primary somatosensory cortex, trunk region | s1tr |
| primary somatosensory cortex, upper lip region | s1ul |
| primary visual cortex | v1 |
| primary visual cortex, binocular area | v1b |
| primary visual cortex, monocular area | v1m |
| red nucleus | r |
| red nucleus, magnocellular part | rmc |
| red nucleus, parvicellular part | rpc |
| reticular nucleus (prethalamus) | rt |
| retroethmoid nucleus | reth |
| retromammillary nucleus | rm |
| retromammillary nucleus, lateral part | rml |
| retromammillary nucleus, medial part | rmm |
| retroparafascicular nucleus | rpf |
| retrorubral field | rrf |
| retrosplenial dysgranular cortex | rsd |
| retrosplenial granular cortex, a region | rsga |
| retrosplenial granular cortex, b region | rsgb |

| | |
|---|---|
| retrosplenial granular cortex, c region | rsgc |
| retrouniens area | rre |
| reuniens thalamic nucleus | re |
| rhomboid thalamic nucleus | rh |
| rostral amygdalopiriform area | rapir |
| rostral interstitial nucleus | ri |
| rostral linear nucleus (midbrain) | rli |
| secondary auditory cortex, dorsal area | aud |
| secondary auditory cortex, ventral area | auv |
| secondary somatosensory cortex | s2 |
| secondary visual cortex, lateral area | v2l |
| secondary visual cortex, mediolateral area | v2ml |
| secondary visual cortex, mediomedial area | v2mm |
| septofimbrial nucleus | sfi |
| subiculum, transition area | str |
| submedius thalamic nucleus | sub |
| substantia innominata, basal part | sib |
| substantia nigra, compact part, dorsal tier | sncd |
| substantia nigra, reticular part | snr |
| superficial gray layer of the superior colliculus | sug |
| supraoculomotor cap | su3c |
| tectospinal trac | ts |
| temporal association cortex | tea |
| thalamus - unspecified | tunsp |
| ventral anterior thalamic nucleus | va |
| ventral anterior thalamic nucleus/ventrolateral thalamic nucleus | va/vl |
| ventral intermediate entorhinal cortex | vient |
| ventral orbital cortex | vo |
| ventral pallidum | vp |
| ventral part of claustrum | vcl |
| ventral posterior nucleus of the thalamus, parvicellular part | vppc |
| ventral posterolateral thalamic nucleus | vpl |
| ventral posteromedial thalamic nucleus | vpm |
| ventral subiculum | vs |
| ventral tegmental area | vta |
| ventrolateral thalamic nucleus | vl |
| ventromedial hypothalamic nucleus | vmh |
| ventromedial thalamic nucleus | vm |
| zona incerta | zi |
| zona incerta, caudal part | zic |
| zona incerta, dorsal part | zid |
| zona incerta, ventral part | ziv |

**Table S2. Statistics of the comparisons made in Figure 3.**

| Comparison | Measure | Test | Significance |
|---|---|---|---|
| Figure 3D: Starter cell analysis, interneurons | % dual-labeled neurons by region | 2way ANOVA; Interaction: F(21,64)=0.90; p=0.58 Region: F(7,64)=60.6; p<0.0001 Cre Line: F(3,64)=0.17; p=0.91 | Tukey's post-hoc tests: p<0.0001 for IL vs. each region in each line |
| Figure 3D: Starter cell analysis, projection neurons | % dual-labeled neurons by region | 2way ANOVA; Interaction: F(21,72)=0.83; p=0.67 Region: F(7,72)=59.92; p<0.0001 Cre Line: F(3,72)=0.006; p=0.99 | Tukey's post-hoc tests: p<0.001 for IL vs. each region in each line |
| Figure 3E: Starter cell correlations, interneurons | Correlation between starter cells and total cells | Two-tailed Spearman n=12 pairs | r= 0.8266, p<0.0001 |
| Figure 3E: Starter cell correlations, projection neurons | Correlation between starter cells and total cells | Two-tailed Spearman n=13 pairs | r= 0.7912, p=0.002 |
| Figure 3E: Starter cell correlations, all | Correlation between starter cells and total cells | Two-tailed Spearman n=25 pairs | r= 0.8058, p<0.0001 |

N.B.- all p-values reflect adjusted values for multiple comparisons

**Table S3. Statistics of the comparisons made in Figure 4.**

| Comparison | Measure | Test | Significance |
|---|---|---|---|
| Figure 4B: Input fraction along anterior-posterior axis, pooled | input fraction at location | 2way ANOVA;<br>Interaction: F(14,360)=1.68; p=0.05<br>Location: F(14,360)=26.03; p<0.0001<br>Cell class: F(1,360)=2e-13; p>0.99 | Sidak's post-hoc test:<br><br>none |
| Figure 4B: Input fraction along anterior-posterior axis, interneurons | input fraction at location | 2way ANOVA;<br>Interaction: F(42,120)=6.40; p<0.0001<br>Location: F(14,120)=28.62; p<0.0001<br>Cell class: F(3,120)=3e-10; p>0.99 | Tukey's post-hoc test:<br>**-0.5 to -1:**<br>PV vs NDNF, p<0.0001<br>SST vs NDNF, p<0.0001<br>VIP vs NDNF, p<0.0001<br>**-1 to -1.5:**<br>PV vs NDNF, p<0.0001<br>SST vs NDNF, p<0.0001<br>VIP vs NDNF, p<0.0001<br>**-3 to -3.5**:<br>PV vs NDNF, p<0.005<br>VIP vs NDNF, p<0.005<br>**-3.5 to -4:**<br>PV vs NDNF, p<0.0001<br>SST vs NDNF, p<0.0001<br>VIP vs NDNF, p<0.0001 |
| Figure 4B: Input fraction along anterior-posterior axis, projections | input fraction at location | 2way ANOVA;<br>Interaction: F(42,150)=1.98; p=0.001<br>Location: F(14,150)=17.0; p<0.0001<br>Cell class: F(3,150)=6e-11; p=>0.99 | Tukey's post-hoc test:<br>**1.5 to 1.0:**<br>BLA vs RE, p=0.03<br>**-0.5 to -1:**<br>LEC vs RE, p=0.003;<br>PAG vs RE, p=0.0004<br>**-3.5 to -4:**<br>BLA vs LEC, p<0.0001;<br>BLA vs PAG, p=0.01;<br>LEC vs RE, p=0.001 |
| Figure 4C: Input fraction at anterior peak, interneurons | input fraction at location | 2way ANOVA;<br>Interaction: F(18,56)=36.58; p<0.0001<br>Region: F(6,56)=276.5; p<0.0001<br>Cell class: F(3,56)=42.89; p<0.0001 | Tukey's post-hoc test:<br>**Thalamus:**<br>PV vs NDNF, p<0.0001<br>SST vs NDNF, p<0.0001<br>VIP vs NDNF, p<0.0001 |
| Figure 4C: Input fraction at anterior peak, projections | input fraction at location | 2way ANOVA;<br>Interaction: F(18,70)=4.85; p<0.0001 | Tukey's post-hoc test:<br>**Thalamus:**<br>BLA vs LEC, p<0.0001;<br>BLA vs. RE, p=0.009;<br>BLA vs PAG, p<0.0001;<br>LEC vs RE, p<0.0001; |

| | | | |
|---|---|---|---|
| | | Region: F(6,70)=93.29; p<0.0001<br>Cell class: F(3,70)=4.41; p=0.006 | RE vs PAG, p<0.0001 |
| Figure 4C: Input fraction at posterior peak, interneurons | input fraction at location | 2way ANOVA;<br>Interaction: F(9,32)=3.94; p=0.001<br>Region: F(9,32)=68.62; p<0.0001<br>Cell class: F(3,32)=3.76; p=0.02 | Tukey's post-hoc test:<br>**Hippocampus:**<br>PV vs NDNF, p<0.0005<br>SST vs NDNF, p<0.0005<br>VIP vs NDNF, p<0.0005 |
| Figure 4C: Input fraction at posterior peak, projections | input fraction at location | 2way ANOVA;<br>Interaction: F(9,40)=1.28; p=0.27<br>Region: F(3,40)=65.46; p<0.0001<br>Cell class: F(3,40)=2.11; p=0.11 | Tukey's post-hoc test:<br>**Hippocampus:**<br>BLA vs LEC, p=0.001;<br>LEC vs RE, p=0.02 |
| Figure 4D: Input fraction across the brain | input fraction across brain | 2way ANOVA;<br>Interaction: F(18,56)=13.51; p<0.0001<br>Region: F(6,56)=145.1; p<0.0001<br>Cell class: F(3,56)=0.35; p=0.78 | Tukey's post-hoc test:<br>**Hippocampus:** PV vs NDNF, p<0.0001<br>SST vs NDNF, p<0.0001<br>VIP vs NDNF, p<0.0001<br>**Thalamus:** PV vs NDNF, p<0.0001<br>SST vs NDNF, p<0.0001<br>VIP vs NDNF, p<0.0001<br>**Cortex:** PV vs VIP, p=0.007;<br>PV vs NDNF, p=0.01;<br>SST vs VIP, p=0.04; |

N.B.- all p-values reflect adjusted values for multiple comparisons

**Table S4. Statistics of the comparisons made in Figure 5.**

| Comparison | Measure | Test | Significance |
|---|---|---|---|
| Figure 5A: Input fraction within thalamus, interneurons | input fraction | 2way ANOVA;<br>Interaction:<br>F(12,40)=7.82;<br>p<0.0001<br>Location:<br>F(4,40)=36.48;<br>p<0.0001<br>Cell class:<br>F(3,40)=19.41;<br>p<0.0001 | Tukey's post-hoc test:<br>**Secondary thalamus:**<br>PV vs NDNF, p<0.0001<br>SST vs NDNF, p<0.0001<br>VIP vs NDNF, p<0.0001<br>**Reuniens:**<br>PV vs NDNF, p<0.0001<br>SST vs NDNF, p<0.0001<br>VIP vs NDNF, p<0.0001 |
| Figure 5A: Input fraction within cortex, interneurons | input fraction | 2way ANOVA;<br>Interaction:<br>F(15,48)=2.16;<br>p=0.02<br>Location:<br>F(5,48)=112.1;<br>p<0.0001<br>Cell class:<br>F(3,48)=3.9; p=0.01 | Tukey's post-hoc test:<br>**Frontal Association:**<br>PV vs VIP, p=0.0006;<br>PV vs NDNF, p=0.0001;<br>SST vs VIP, p=0.01;<br>SST vs NDNF, p=0.002 |
| Figure 5B: Input fraction at individual regions | input fraction at location | 2way ANOVA;<br>Interaction:<br>F(633,1696)=5.7;<br>p<0.0001<br>Location:<br>F(211,1696)=57.8;<br>p<0.0001<br>Cell class:<br>F(3,1696)=0.43;<br>p=0.72 | Tukey's post-hoc test:<br>**BLA:** SST vs. NDNF, p=0.0009<br>**BLP:** PV vs. NDNF, p=0.006<br>**AI:** PV vs SST, p<0.0001;<br>PV vs NDNF, p=0.009;<br>SST vs VIP, p<0.0001;<br>SST vs NDNF, p<0.0001<br>**CG1/ACC:**<br>PV vs SST, p<0.0001;<br>PV vs VIP, p<0.0001;<br>PV vs NDNF, p<0.0001<br>**CG2/PL:**<br>PV vs SST, p=0.04<br>**M2:**<br>PV vs VIP, p=0.01<br>PV vs NDNF, p<0.0001;<br>SST vs NDNF, p=0.0004<br>**PIR:**<br>PV vs SST, p<0.0001;<br>PV vs VIP, p<0.0001;<br>PV vs NDNF, p<0.0001;<br>SST vs NDNF, p<0.0001;<br>VIP vs NDNF, p<0.0001<br>**PL:**<br>PV vs SST, p=0.02;<br>PV vs VIP, p=0.02;<br>**S2:**<br>PV vs NDNF, p=0.003;<br>SST vs NDNF, p<0.0001;<br>VIP vs NDNF, p=0.0006<br>**CA1:**<br>PV vs SST, p<0.0001;<br>SST vs VIP, P<0.0001; |

| | | | |
|---|---|---|---|
| | | | PV vs NDNF, p<0.0001<br>SST vs NDNF, p<0.0001<br>VIP vs NDNF, p<0.0001<br>**DLENT:**<br>SST vs NDNF, p=0.0009<br>**DS:**<br>PV/SST/VIP vs NDNF, p<0.0001<br>**STR:**<br>PV vs VIP, p=0.01;<br>PV vs NDNF, p<0.0001;<br>SST vs NDNF, p<0.0001<br>VIP vs NDNF, p<0.0001<br>**CLAUS:**<br>PV vs NDNF, p=0.0001<br>SST vs NDNF, p=0.001<br>**INS:**<br>PV vs SST, p=0.003;<br>SST vs VIP, p=0.0007;<br>SST vs NDNF, p=0.009<br>**OLF:**<br>VIP vs NDNF, p=0.03<br>**AM:**<br>PV vs NDNF, p<0.0001<br>SST vs NDNF, p<0.0001<br>VIP vs NDNF, p<0.0001<br>SST vs VIP, p=0.002<br>**MD:**<br>PV vs NDNF, p=0.0002<br>SST vs NDNF, p=0.0007<br>VIP vs NDNF, p=0.0001<br>**PV:**<br>SST vs NDNF, p=0.008<br>**RE:**<br>PV vs NDNF, p<0.0001<br>SST vs NDNF, p<0.0001<br>VIP vs NDNF, p<0.0001<br>**SUB:**<br>PV vs NDNF, p<0.0001<br>SST vs NDNF, p<0.0001<br>VIP vs NDNF, p<0.0001<br>**VA/VL:**<br>SST/VIP vs NDNF, p=0.04<br>**VM:**<br>PV vs NDNF, p<0.0001<br>SST vs NDNF, p<0.0001<br>VIP vs NDNF, p<0.0001 |

N.B.- all p-values reflect adjusted values for multiple comparisons

**Table S5. Statistics of the comparisons made in Figure 6.**

| Comparison | Measure | Test | Significance |
|---|---|---|---|
| Figure 6A: Input fraction within thalamus, projection neurons | input fraction | 2way ANOVA; Interaction: F(12,50)=3.58, p=0.0007 Location: F(4,50)=42.88; p<0.0001 Cell class: F(3,50)=10.61; p<0.0001 | Tukey's post-hoc test: **Secondary thalamus:** BLA vs LEC, p=0.002; BLA vs RE, p=0.02; BLA vs PAG, p<0.0001; LEC vs RE, p<0.0001; RE vs PAG, p<0.0001 |
| Figure 6A: Input fraction within cortex, projection neurons | input fraction | 2way ANOVA; Interaction: F(15,60)=2.88; p=0.001 Location: F(5,60)=63.26; p<0.0001 Cell class: F(3,60)=0.27; p=0.84 | Tukey's post-hoc test: **Frontal Association:** BLA vs LEC, p=0.005; LEC vs RE, p=0.01; **Temporal Association:** BLA vs LEC, p=0.0002 LEC vs RE, p=0.01 LEC vs PAG, p=0.03 |
| Figure 6B: Input fraction at individual regions | input fraction at location | 2way ANOVA; Interaction: F(633, 2120)=1.68; p<0.0001 Location: F(211, 2120)=49.45; p<0.0001 Cell class: F(3,2120)=0.13; p=0.93 | Tukey's post-hoc test: **BLA:** BLA vs PAG, p=0.01; RE vs PAG, p=0.01 **BLP:** LEC vs. RE, p=0.02; RE vs. PAG, p=0.01 **CG1/ACC:** BLA vs PAG, p=0.0005; LEC vs PAG, p=0.04; **CG2:** BLA vs PAG, p=0.008 **IL**: BLA vs RE, p=0.01 **LO**: BLA vs LEC, p=0.04 **M2**: BLA vs LEC, p<0.0001 LEC vs RE, p=0.01; LEC vs PAG, p=0.02 **PIR**: BLA vs LEC, p<0.0001; BLA vs RE, p=0.02 BLA vs PAG, p<0.0001 LEC vs RE, p<0.0001 RE vs PAG, p<0.0001 **PL:** BLA vs LEC, p=0.0005; LEC vs RE, p=0.0007; LEC vs PAG, p=0.04 **CA1:** BLA vs LEC, p=0.0002; BLA vs PAG, p<0.0001; **DLENT:** BLA vs LEC, p<0.0001 |

| | | | |
|---|---|---|---|
| | | | BLA vs PAG, p=0.02;<br>LEC vs RE, p=0.0004<br>**DS:**<br>BLA vs LEC, p=0.001<br>BLA vs RE, p<0.0001;<br>LEC vs RE, p<0.0001;<br>LEC vs PAG, p=0.02;<br>RE vs PAG, p<0.0001<br>**STR:**<br>BLA vs LEC, p<0.0001;<br>LEC vs RE, p<0.0001;<br>LEC vs PAG, p<0.0001<br>**VS:**<br>BLA vs LEC, p=0.0002<br>BLA vs RE, p<0.0001<br>BLA vs PAG, p=0.0002<br>**DP:**<br>BLA vs RE, p<0.0001<br>LAC vs RE, p<0.0001<br>RE vs PAG, p>0.0001<br>**INS:**<br>BLA vs LEC, p=0.002;<br>BLA vs PAG, p=0.03;<br>**LS:**<br>BLA vs RE, p<0.0001<br>LAC vs RE, p<0.0001<br>RE vs PAG, p<0.0001<br>**OLF:**<br>BLA vs RE, p=0.04;<br>RE vs PAG, p=0.02<br>**PRH:**<br>BLA vs LEC, p=0.03<br>**AM:**<br>BLA vs LEC, p<0.0001;<br>BLA vs PAG, p<0.0001;<br>LEC vs RE, p<0.0001;<br>LEC vs PAG, p=0.006<br>RE vs PAG, p<0.0001<br>**MD:**<br>BLA vs LEC, p=0.0003<br>LEC vs RE, p<0.0001<br>LEC vs PAG, p=0.001<br>**MDL:**<br>BLA vs PAG, p=0.04<br>RE vs PAG, p=0.005<br>**PV:**<br>BLA vs RE, p=0.01<br>LEC vs RE, p=0.002<br>LEC vs PAG, p=0.04<br>**RE:**<br>LEC vs RE, p=0.01;<br>**VMT:**<br>RE vs PAG, p=0.03; |

N.B.- all p-values reflect adjusted values for multiple comparisons

**Table S6. Statistics of the comparisons made in Supplementary Figure 2.**

| Comparison | Measure | Test | Significance |
|---|---|---|---|
| Supplementary Figure 2B: interneuron dendritic morphology | dendritic material by distance from soma | 2way ANOVA;<br>Interaction: F(63,506)=6.13; p<0.0001<br>Distance: F(21,506)=87.76; p<0.0001<br>Cre Line: F(3,506)=75.84; p<0.0001 | Tukey's post-hoc tests:<br>**0-20µm:**<br>PV vs. VIP, p=0.001;<br>VIP vs NDNF, p=0.009<br>**20-40µm:**<br>PV vs VIP, p=0.0001<br>PV vs NDNF, p=0.03<br>VIP vs NDNF, p<0.0001<br>**40-60µm:**<br>PV vs VIP, p=0.0004<br>SST vs VIP, p=0.01<br>VIP vs NDNF, p=0.0008<br>**60-80µm:**<br>PV vs VIP, p=0.04<br>PV vs NDNF, p=0.003<br>SST vs VIP, p=0.0006<br>SST vs NDNF, p<0.0001<br>**80-100µm:**<br>PV vs SST, p=0.01<br>PV vs NDNF, p<0.0001<br>SST vs VIP, p=0.0006<br>SST vs NDNF, p<0.0001<br>VIP vs NDNF, p=0.0006<br>**100-120µm:**<br>PV vs SST, p=0.004<br>PV vs NDNF, p<0.0001<br>SST vs VIP, p=0.003<br>SST vs. NDNF, p<0.0001<br>VIP vs NDNF, p<0.0001<br>**120-140µm:**<br>PV vs SST, p=0.0002<br>PV vs NDNF, p<0.0001<br>SST vs VIP, p=0.0006<br>SST vs NDNF, p<0.0001<br>VIP vs NDNF, p<0.0001<br>**140-160µm:**<br>PV vs SST, p=0.0004<br>PV vs NDNF, p<0.0001<br>SST vs VIP, p=0.003<br>SST vs NDNF, p<0.0001<br>VIP vs NDNF, p<0.0001<br>160-180µm:<br>PV vs SST, p=0.002<br>PV vs NDNF, p<0.0001<br>SST vs VIP, p=0.04<br>SST vs NDNF, p<0.0001<br>VIP vs NDNF, p<0.0001<br>180-200µm:<br>PV vs NDNF, p<0.0001<br>SST vs NDNF, p<0.0001<br>VIP vs NDNF, p<0.0001 |

| | | | |
|---|---|---|---|
| | | | 200-220µm:<br>PV vs NDNF, p=0.0004<br>SST vs NDNF, p=0.004<br>VIP vs NDNF, p<0.0001<br>220-240µm:<br>PV vs NDNF, p=0.02<br>240-260µm:<br>SST vs NDNF, p=0.04 |
| Supplementary Figure 2D: projection neuron dendritic morphology | dendritic material by distance from soma | 2way ANOVA;<br>Interaction: F(129,792)=2.69; p<0.0001<br>Distance from Soma: F(43,792)=31.15; p<0.0001<br>Projection: F(3,792)=7.09; p=0.0001 | Tukey's post-hoc tests:<br>Basal: none<br>Apical: 20-40 µm:<br>RE vs LEC, p=0.009;<br>PAG vs. LEC, p=0.02<br>40-60 µm:<br>RE vs LEC, p=0.01;<br>PAG vs. LEC, p=0.01<br>60-80 µm:<br>RE vs LEC, p=0.03;<br>PAG vs. LEC, p=0.001;<br>80-100 µm:<br>RE vs LEC, p=0.003;<br>PAG vs. LEC, p=0.006<br>140-160 µm:<br>PAG vs. LEC, p=0.03<br>320-340 µm:<br>RE vs BLA, p=0.02;<br>RE vs. LEC, p=0.02<br>340-360 µm:<br>RE vs BLA, p=0.01;<br>RE vs. LEC, p=0.002<br>PAG vs LEC, p=0.01<br>360-380 µm:<br>RE vs BLA, p=0.04;<br>RE vs. LEC, p=0.0004<br>PAG vs BLA, p=0.02<br>PAG vs LEC, p=0.0003<br>380-400 µm:<br>RE vs BLA, p=0.0003;<br>RE vs. LEC, p<0.0001<br>PAG vs BLA, p=0.0002<br>PAG vs LEC, p<0.0001<br>400-420 µm:<br>RE vs BLA, p=0.01;<br>RE vs. LEC, p=0.004<br>PAG vs BLA, p<0.0001<br>PAG vs LEC, p<0.0001<br>420-440 µm:<br>RE vs BLA, p=0.03<br>RE vs. LEC, p=0.01<br>PAG vs BLA, p<0.0001<br>PAG vs LEC, p<0.0001<br>440-460 µm:<br>RE vs PAG, p=0.03<br>RE vs BLA, p=0.02; |

| | | | |
|---|---|---|---|
| | | | RE vs. LEC, p=0.007<br>PAG vs BLA, p<0.0001<br>PAG vs LEC, p<0.0001<br>460-480 µm:<br>RE vs BLA, p=0.02;<br>RE vs. LEC, p=0.03<br>PAG vs BLA, p=0.0003<br>PAG vs LEC, p=0.0006<br>480-500 µm:<br>RE vs BLA, p=0.03 |

N.B.- all p-values reflect adjusted values for multiple comparisons

**Table S7. Statistics of the comparisons made in Supplementary Figure 3.**

| Comparison | Measure | Test | Significance |
| --- | --- | --- | --- |
| Supplementary Figure 3B: automatic vs manual counts | Correlations between cell numbers | Two-tailed Spearman n=9 region pairs | r=0.95, p<0.0004 |
| Supplementary Figure 3B: adjacent section counts | Correlations between cell numbers | Two-tailed Spearman n=91 region pairs | r=0.89, p<0.0001 |
| Supplementary Figure 3C: GFP+ neurons from datasets with no Cre vs with Cre | Cell number | Welch-corrected two tailed t-test: df=25.02, t=7.245 | Cre transgene vs. no cre, p<0.0001 |

**Table S8. Statistics of the comparisons made in Supplementary Figure 5.**

| Comparison | Measure | Test | Significance |
|---|---|---|---|
| Supplementary Figure 5C: Correlations among Cre lines | Pairwise correlations in input fraction | Two-tailed Spearman n=32 region pairs | PV vs SST, r=0.74, p<0.0001<br>PV vs VIP, r=0.82, p<0.0001<br>PV vs NDNF, r=0.22, p=0.22<br>SST vs VIP, r=0.75, p<0.0001<br>SST vs NDNF, r=0.19, p=0.29<br>VIP vs NDNF, r=0.35, p=0.04 |

**Table S9. Statistics of the comparisons made in Supplementary Figure 6.**

| Comparison | Measure | Test | Significance |
|---|---|---|---|
| Supplementary Figure 6C: Correlations among projection neurons | Pairwise correlations in input fraction | Two-tailed Spearman n=38 region pairs | BLA vs LEC, r=0.41, p=0.01<br>BLA vs RE, r=0.31, p=0.05<br>BLA vs PAG, r=0.48, p=0.002<br>LEC vs RE, r=0.19, p=0.24<br>LEC vs PAG, r=0.51, p=0.0009<br>RE vs PAG, r=0.27, p=0.09 |

**Table S10. Statistics of the comparisons made in Supplementary Figure 7.**

| Comparison | Measure | Test | Significance |
|---|---|---|---|
| Supplementary Figure 7: Full Two-way ANOVA among all IL postsynaptic cell classes | input fraction at location | 2way ANOVA; Interaction: F(1477, 3816)=3.36; p<0.0001 Location: F(211, 3816)=101.2; p<0.0001 Cell class: F(7, 3816)=0.23; p=0.97 | Tukey's post-hoc tests: **AI:** PV vs. SST, p<0.0001; PV vs. RE, p=0.0003; PV vs. PAG, p=0.0003; SST vs. VIP, p<0.0001; SST vs. NDNF, p<0.0001; SST vs. BLA, p<0.0001; SST vs. LEC, p<0.0001; SST vs. RE, p<0.0001; SST vs. PAG, p<0.0001 **AM**: PV vs. NDNF, p<0.0001; PV vs. LEC, p<0.0001; PV vs. PAG, p<0.0001; SST vs. VIP, p=0.02; SST vs. NDNF, p<0.0001; SST vs. BLA, p<0.0001; SST vs. LEC, p<0.0001; SST vs. RE, p<0.0001; SST vs. PAG, p<0.0001 VIP vs. NDNF, p<0.0001; VIP vs. RE, p=0.009; VIP vs. PAG, p<0.0001; NDNF vs. BLA, p<0.0001; NDNF vs. LEC, p<0.0001; NDNF vs. RE, p<0.0001; BLA vs. LEC, p<0.0001; BLA vs. PAG, p<0.0001; LEC vs. RE, p<0.0001; LEC vs. PAG, p=0.01; RE vs. PAG, p<0.0001 **BLA, anterior part:** PV vs. PAG, p=0.002; SST vs. NDNF, p=0.01; VIP vs. PAG, p=0.005; NDNF vs. BLA, p=0.01; NDNF vs. LEC, p=0.01; NDNF vs. PAG, p<0.0001; BLA vs. PAG, p=0.04; RE vs. PAG, p=0.02 **BLA, posterior part:** NDNF vs. LEC, p=0.02; NDNF vs. PAG, p=0.01; RE vs. PAG, p=0.03 **CA1:** PV vs. SST, p<0.0001; PV vs. NDNF, p<0.0001; PV vs. LEC, p<0.0001; PV vs. RE, p=0.002; PV vs. PAG, p<0.0001; SST vs. VIP, p<0.0001; SST vs. NDNF, p<0.0001; SST vs. BLA, p<0.0001; SST vs. LEC, p=0.007; |

| | | | |
|---|---|---|---|
| | | | SST vs. RE, p<0.0001;<br>SST vs. PAG, p=0.03;<br>VIP vs. NDNF, p<0.0001;<br>VIP vs. LEC, p=0.001;<br>VIP vs. PAG, p=0.0001;<br>NDNF vs. BLA, p<0.0001;<br>NDNF vs. LEC, p<0.0001;<br>NDNF vs. RE, p<0.0001;<br>NDNF vs. PAG, p<0.0001;<br>BLA vs. LEC, p=0.0004;<br>BLA vs. PAG, p<0.0001<br>**CG1/ACC:**<br>PV vs. SST, p=0.0009;<br>PV vs. VIP, p=0.0001;<br>PV vs. NDNF, p<0.0001;<br>PV vs. BLA, p<0.0001;<br>PV vs. LEC, p<0.0001;<br>PV vs. RE, p<0.0001;<br>NDNF vs. PAG, p=0.006;<br>BLA vs. PAG, p=0.0009<br>**CG2:**<br>SST vs. PAG, p=0.03;<br>NDNF vs. PAG, p=0.001;<br>BLA vs. PAG, p=0.02<br>**CLAUS:**<br>PV vs. NDNF, p=0.001;<br>SST vs. NDNF, p=0.01<br>**DLEC:**<br>PV vs. LEC, p=0.004;<br>SST vs. NDNF, p=0.01;<br>VIP vs. LEC, p=0.003;<br>NDNF vs. LEC, p<0.0001;<br>NDNF vs. PAG, p=0.008;<br>BLA vs. LEC, p<0.0001;<br>LEC vs. RE, p=0.0008<br>**DP:**<br>PV vs. RE, p=0.0001;<br>SST vs. RE, p<0.0001;<br>VIP vs. RE, p<0.0001;<br>NDNF vs. RE, p<0.0001;<br>BLA vs. RE, p<0.0001;<br>LEC vs. RE, p<0.0001;<br>RE vs. PAG, p<0.0001<br>**DS:**<br>PV vs. NDNF, p<0.0001;<br>PV vs. LEC, p<0.0001;<br>PV vs. RE, p<0.0001;<br>SST vs. NDNF, p<0.0001;<br>SST vs. BLA, p=0.005;<br>SST vs. LEC, p<0.0001;<br>SST vs. RE, p<0.0001;<br>SST vs. PAG, p=0.003;<br>VIP vs. NDNF, p<0.0001;<br>VIP vs. LEC, p<0.0001;<br>VIP vs. RE, p<0.0001; |

| | | | |
|---|---|---|---|
| | | | VIP vs. PAG, p=0.03;<br>NDNF vs. BLA, p<0.0001;<br>NDNF vs. RE, p<0.0001;<br>NDNF vs. PAG, p<0.0001;<br>BLA vs. LEC, p=0.003;<br>BLA vs. RE, p<0.0001;<br>LEC vs. RE, p<0.0001;<br>RE vs. PAG, p<0.0001<br>**IL:**<br>SST vs. RE, p=0.04;<br>VIP vs. RE, p=0.02;<br>BLA vs. RE, p=0.03<br>**INS:**<br>PV vs. SST, p=0.03;<br>SST vs. VIP, p=0.008;<br>SST vs. LEC, p<0.0001;<br>SST vs. RE, p=0.0001;<br>SST vs. PAG, p<0.0001;<br>BLA vs. LEC, p=0.004<br>**LS:**<br>PV vs. RE, p=0.005;<br>SST vs. RE, p<0.0001;<br>VIP vs. RE, p=0.003;<br>NDNF vs. RE, p<0.0001;<br>BLA vs. RE, p<0.0001;<br>LEC vs. RE, p=0.0001;<br>RE vs. PAG, p<0.0001<br>**M2:**<br>PV vs. NDNF, p<0.0001;<br>PV vs. BLA, p<0.0001;<br>SST vs. NDNF, p=0.005;<br>SST vs. BLA, p=0.001;<br>VIP vs. LEC, p=0.03;<br>NDNF vs. LEC, p<0.0001;<br>BLA vs. LEC, p<0.0001;<br>LEC vs. RE, p=0.03;<br>LEC vs. PAG, p=0.04<br>**MD:**<br>PV vs. NDNF, p=0.003;<br>PV vs. LEC, p<0.0001;<br>SST vs. NDNF, p=0.008;<br>SST vs. LEC, p<0.0001;<br>VIP vs. NDNF, p=0.002;<br>VIP vs. LEC, p<0.0001;<br>NDNF vs. RE, p=0.0004;<br>BLA vs. LEC, p=0.0005;<br>LEC vs. RE, p<0.0001;<br>LEC vs. PAG, p=0.002<br>**MDL:**<br>RE vs. PAG, p=0.01<br>**PIR:**<br>PV vs. SST, p=0.0005;<br>PV vs. VIP, p<0.0001;<br>PV vs. NDNF, p<0.0001;<br>PV vs. LEC, p<0.0001; |

| | | | |
|---|---|---|---|
| | | | PV vs. RE, p=0.005;<br>PV vs. PAG, p<0.0001;<br>SST vs. NDNF, p<0.0001;<br>SST vs. BLA, p=0.008;<br>SST vs. LEC, p<0.0001;<br>SST vs. PAG, p<0.0001;<br>VIP vs. NDNF, p<0.0001;<br>VIP vs. BLA, p<0.0001;<br>VIP vs. LEC, p<0.0001;<br>VIP vs. PAG, p<0.0001;<br>NDNF vs. BLA, p<0.0001;<br>NDNF vs. LEC, p<0.0001;<br>NDNF vs. RE, p<0.0001;<br>NDNF vs. PAG, p<0.0001;<br>BLA vs. LEC, p<0.0001;<br>BLA vs. PAG, p<0.0001;<br>LEC vs. RE, p<0.0001;<br>RE vs. PAG, p<0.0001<br>**PL:**<br>PV vs. BLA, p=0.01;<br>PV vs. RE, p=0.01;<br>SST vs. LEC, p=0.01;<br>VIP vs. LEC, p=0.01;<br>BLA vs. LEC, p=0.0009;<br>LEC vs. RE, p=0.001<br>**PRH:**<br>PV vs. LEC, p=0.001;<br>PV vs. PAG, p=0.03;<br>SST vs. LEC, p=0.006;<br>NDNF vs. LEC, p=0.007<br>**PVT:**<br>PV vs. BLA, p=0.02;<br>PV vs. LEC, p=0.003;<br>VIP vs. BLA, p=0.01;<br>VIP vs. LEC, p=0.001;<br>NDNF vs. BLA, p=0.0001;<br>NDNF vs. LEC, p<0.0001;<br>BLA vs. RE, p=0.02;<br>LEC vs. RE, p=0.004<br>**RE:**<br>PV vs. NDNF, p<0.0001;<br>PV vs. BLA, p<0.0001;<br>PV vs. LEC, p<0.0001;<br>PV vs. RE, p=0.04;<br>PV vs. PAG, p=0.0006;<br>SST vs. NDNF, p<0.0001;<br>SST vs. BLA, p=0.004;<br>SST vs. LEC, p<0.0001;<br>SST vs. PAG, p=0.01;<br>VIP vs. NDNF, p<0.0001;<br>VIP vs. LEC, p=0.0001;<br>NDNF vs. BLA, p<0.0001;<br>NDNF vs. LEC, p<0.0001;<br>NDNF vs. RE, p<0.0001;<br>NDNF vs. PAG, p<0.0001; |

| | | | |
|---|---|---|---|
| | | | LEC vs. RE, p=0.03<br>**S2:**<br>PV vs. NDNF, p=0.03;<br>SST vs. NDNF, p=0.001;<br>VIP vs. NDNF, p=0.007;<br>NDNF vs. BLA, p=0.0002;<br>NDNF vs. LEC, p=0.002;<br>NDNF vs. RE, p=0.009;<br>NDNF vs. PAG, p=0.002<br>**STR:**<br>PV vs. NDNF, p<0.0001;<br>PV vs. LEC, p<0.0001;<br>SST vs. NDNF, p<0.0001;<br>SST vs. BLA, p=0.009;<br>SST vs. LEC, p<0.0001;<br>SST vs. PAG, p=0.01;<br>VIP vs. NDNF, p<0.0001;<br>VIP vs. BLA, p<0.0001;<br>VIP vs. LEC, p=0.001;<br>VIP vs. RE, p=0.004;<br>VIP vs. PAG, p<0.0001;<br>NDNF vs. BLA, p<0.0001;<br>NDNF vs. RE, p<0.0001;<br>NDNF vs. PAG, p<0.0001;<br>BLA vs. LEC, p<0.0001;<br>LEC vs. RE, p<0.0001;<br>LEC vs. PAG, p<0.0001<br>**SUB:**<br>PV vs. NDNF, p=0.0002;<br>SST vs. NDNF, p=0.0002;<br>VIP vs. NDNF, p=0.001;<br>NDNF vs. BLA, p<0.0001;<br>NDNF vs. LEC, p=0.0007;<br>NDNF vs. RE, p<0.0001;<br>NDNF vs. PAG, p<0.0001<br>**VM:**<br>PV vs. NDNF, p<0.0001;<br>SST vs. NDNF, p<0.0001;<br>VIP vs. NDNF, p<0.0001;<br>NDNF vs. BLA, p<0.0001;<br>NDNF vs. LEC, p<0.0001;<br>NDNF vs. RE, p<0.0001;<br>NDNF vs. PAG, p<0.0001<br>**VS:**<br>PV vs. BLA, p<0.0001;<br>SST vs. BLA, p<0.0001;<br>VIP vs. BLA, p=0.0003;<br>NDNF vs. BLA, p<0.0001;<br>BLA vs. LEC, p=0.0003;<br>BLA vs. RE, p<0.0001;<br>BLA vs. PAG, p=0.0004 |

N.B.- all p-values reflect adjusted values for multiple comparisons
